# A live cell biosensor protocol for high-resolution screening of therapy-resistant cancer cells

**DOI:** 10.1101/2025.09.23.678038

**Authors:** Viral D. Oza, Colin Williams, Jessica S. Blackburn

## Abstract

The Genetically Encoded Death Indicator (GEDI) is a ratiometric, dual-fluorescence biosensor that enables real-time detection of cell death through calcium influx. Originally developed for use in neurodegeneration models, GEDI can be applied to cancer cells to quantify therapy-induced death at single-cell resolution. This protocol details how to generate GEDI-expressing cancer cell lines, empirically determine stress-induced GEDI thresholds using radiation or chemotherapeutic agents, and perform time-resolved imaging and image analysis to track cell fate. This workflow is optimized for high-throughput drug and radiation screening in heterogeneous populations and is especially useful for identifying chemo- and radio-resistant subclones. Key limitations include the need for empirical GEDI threshold calibration for each treatment condition and careful standardization of imaging parameters. The protocol outputs include GEDI ratio values, single-cell time-of-death annotations, and whole-cell morphological data in parallel, which can be linked to downstream applications such as FACS-based isolation of live or dying subpopulations, transcriptomic profiling of resistant clones, or in vivo validation using xenografts or organotypic slice culture.

**Associated content:** io.protocol DOI: dx.doi.org/10.17504/protocols.io.eq2ly4d7qlx9/v1

## Introduction

Evasion of apoptosis is a hallmark of cancer and measuring apoptotic events is a key determinant of treatment response^1,2,3^. Programmed cell death can occur as early as 30 minutes after therapy exposure and is influenced by multiple intrinsic and extrinsic pathways, making precise measurement of cell death essential for identifying new vulnerabilities in cancer cells^4,5^. Accurately capturing these events is especially important in drug and radiation screening, where the ability to quantify therapy-induced death is critical for both evaluating treatment efficacy and for detecting resistant subpopulations within heterogeneous tumors.

The heterogeneity within cancer cell lines is at odds with current high-throughput screening (HTS) methods, which typically rely on bulk population readouts of cell death^6^. Within a single population, individual cells may vary widely in their responses, with some succumbing quickly to therapy while others survive due to intrinsic mutations or adaptive stress responses^7,8^. Such heterogeneity has emerged as a major hurdle in cancer treatment, as resistant subclones can evade therapy and ultimately drive relapse. Bulk assays that average outcomes across populations often mask these minority populations, obscuring important information about treatment failure. A method to track cell death with single-cell resolution and precise timing is needed to fully capture how different cells in a heterogeneous tumor respond to therapy. However, conventional cell viability and cytotoxicity assays have significant limitations in this regard. Most HTS assays use static timepoints (24-72 hours) that miss early or transient events. Traditional luminescence-based assays like ATP or caspase activity reporters require cell lysis, providing only a snapshot of cell death and losing spatial or temporal context. Annexin V detection can distinguish apoptotic and dead cells, but requires dyes, flow cytometry, or microscopy at fixed time points, making it less compatible for continuous monitoring. Collectively, these traditional methods lack the ability to monitor cell death dynamically in live, individual cells, making it challenging to resolve mixed responses or rare populations. To overcome these limitations, new tools are needed that combine the throughput of population-based assays with the resolution of live-cell, single-cell imaging.

The Genetically Encoded Death Indicator (GEDI) directly addresses these challenges. GEDI is a dual-fluorescence ratiometric reporter initially developed in neurodegeneration models to measure cell death based on calcium influx^9^. In its original form, GEDI links a constitutive fluorescent reporter (mApple) with a calcium sensor (GC150) through a self-cleaving peptide (P2A) (Figure 1A). GC150 is activated upon an unregulated Ca^2+^ influx, indicating a disruption in membrane integrity (Figure 1B), while mApple provides a whole-cell morphological reference. Normalizing the fluorescence intensity of GC150 to mApple provides a “GEDI ratio”, which serves as an absolute per-cell measure of Ca^2+^ dysregulation that provides a quantitative marker of cell death at single-cell resolution (Figure 1C).

**Figure 1:**
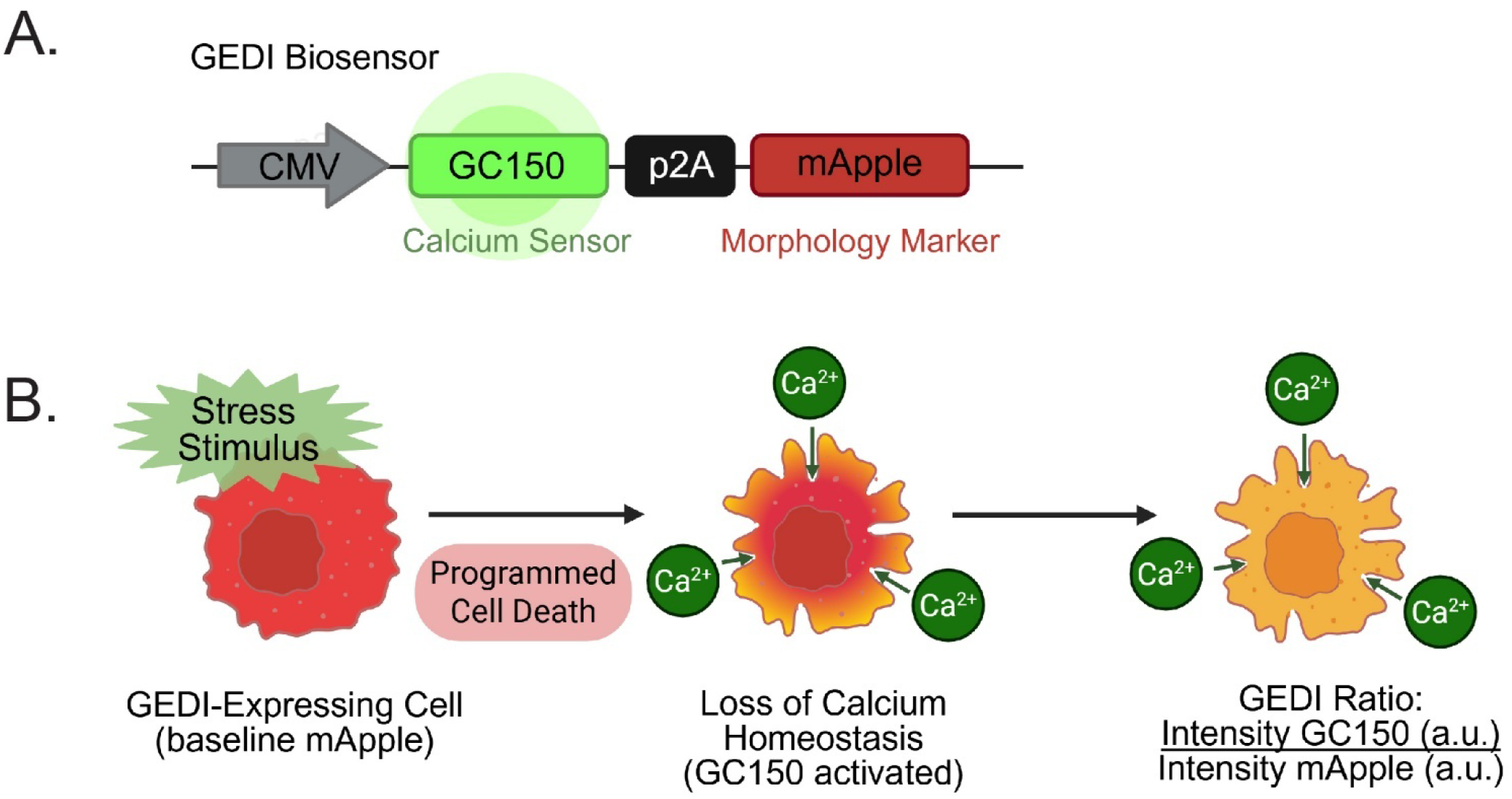
Schematic of the Genetically Encoded Death Indicator (GEDI) biosensor and detection mechanism. (**A**) The GEDI biosensor construct links a calcium sensor (GC150) and a constitutively expressed mApple via a P2A peptide under a CMV promoter. (**B**) The mechanism by which the GEDI biosensor records cell death. A baseline mApple signal provides a morphological reference of a GEDI-expressing cell. Upon a stress stimulus, programmed cell death leads to loss of membrane integrity and increased calcium influx, activating GC150 fluorescence. The GEDI ratio, defined as GC150 fluorescence intensity/mApple fluorescence intensity in arbitrary units (a.u.), provides a quantitative single-cell measure of calcium dysregulation and cell death.

Here, we present a protocol that uses the GEDI biosensor to track therapy-induced death in heterogeneous cancer cell populations. The workflow describes how to generate stable GEDI-expressing lines, empirically determine the GEDI threshold for radiation or drug-induced cell death and perform time-resolved imaging of GEDI-expressing cells with automated cell tracking. By capturing longitudinal phenotypic data in conjunction with real-time cell death events, this approach enables deeper insights into dynamic resistance, clonal evolution, and bystander effects. The protocol is designed for high-throughput drug and radiation screens and supports downstream applications where additional investigations on resistant populations are required, such as *in vivo* or *ex vivo* validation, next-generation sequencing analyses, or cell-specific marker identification.

## Materials and Methods

The protocol described in this peer-reviewed article is published on protocols.io <u>DOI: dx.doi.org/10.17504/protocols.io.eq2ly4d7qlx9/v1</u>and is included for printing as **Supplemental File S1** with this article.

### Cell Culture

H3K27M-pDMG cell line SF8628 (SCC127, Millipore Sigma) was cultured in DMEM-High Glucose (D6546, Sigma) with 10% FBS (S11150H, R&D Systems). Cells were routinely tested for mycoplasma with LookOut Mycoplasma PCR Detection Kit (MP0035, Sigma) and used within 10 passages.

For sub culturing, cells were lifted with Trypsin-EDTA (25300062, Life Technologies) for 5 minutes at 37°C then quenched with 2x volume of media. Cells were then pelleted at 300 x g for 5 minutes, then resuspended in cell culture media. All cells were maintained at 37°C with 5% CO_2_. Media was replaced every 3-4 days and cells were split when confluency reached ∼70-80%.

### Gateway Cloning

To generate an SF8628 GEDI reporter line, the pMe:GC150-p2A-mApple construct (gift from the Finkbeiner Lab, Gladstone Institutes) was cloned into a pLenti-CMV-Puro destination vector (17452, Addgene) using Invitrogen Gateway LR Clonase II (11791020, Life Technologies). The plasmid map can be found in **Supplemental File S2**. Lentivirus was produced in HEK293T cells (CRL-3216, ATCC) via co-transfection with psPAX2 (12260, Addgene) and pMD2.g (12259, Addgene) using TransIT reagent (2300, Mirus Bio). Viral supernatants were collected at 48-72 h, concentrated with Lenti-X (631231, Takara Bio), and used to transduce SF8628 cells. Positive populations were enriched by FACS based on mApple and GC150 expression.

### X-RAY Irradiation

Cells in 96-well plates were irradiated using a XRAD 225XL (PXI Precision X-ray) irradiator with the following beam parameters: 13.3mA, SSD = 40, 225kV, 2.2Gy/min. The irradiation plate was set to a fixed rotation speed. Typical exposure time for 8 Gy was ∼3 minutes. All 0 Gy plates were sham irradiated. Volume of media in culture plates was kept at a total volume of 100 µL.

### JQ1 Preparation

JQ1 (S7110, Selleck Chemicals) was dissolved in DMSO (D8418, Millipore) to 50 mM, followed by a dilution to 10 mM in 200 Proof Ethanol (BP2818-4, Fisher BioReagents) to account for solubility, and used at the indicated concentrations.

### Endpoint Viability Assays

SF8628 cells were plated at a density of 5,000 cells/well in 96-well optical plates (29444, VWR) with 100 µL of media. Cell viability was measured with Caspase-Glo 3/7 (PAG8091, VWR) or Cell Titer-Glo (PAG7572, VWR) according to the manufacturer’s protocol. Luminescence was measured using a Synergy LX Multi-Mode Reader (Biotek). Media-only wells were used as blanks for normalization.

### GEDI Image Analysis

Time-lapse imaging GEDI-expressing cells was conducted using a Lionheart FX imager equipped with red and green filter sets (Biotek), capturing 10 x 10 montages of wells in 96-well plates (29444-008, VWR). Optimal parameters were set at 20X magnification with mApple exposure at 150 ms, and GC150 at 300 ms. Images were processed using Gen5 v3.11 software (Biotek) for background correction, deconvolution, and stitching. GEDI ratios (GC150/mApple) were quantified using ImageJ 1.54p scripts, and empirical thresholds for cell death were determined from high-dose radiation or drug treatment. GEDI ratios were plotted at 0, 3, 6, 12, 18, 24, 36 hours post-stress using the tidyverse package in RStudio. A Rmarkdown file can be found in **S1** detailing how to prepare GEDI data for visualization from ImageJ-generated data.

### Cell Tracking

Automated tracking was performed in ImageJ using TrackMate v7.14^8^ with loG detection and LAP Tracker. GEDI ratios were calculated along tracks as described in **S1**.

### Statistics

Data are presented as mean +/-standard error (SEM) or +/-95% confidence interval where noted. Statistical analyses were performed using Prism 8 (Graphpad software), R (4.3.1), R tidyverse package (2.0.0), and Trackmatev7.14. An annotated ImageJ script and Rstudio markdown file can be found in the io.protocol (**S1**). All experiments had at least 2 biological replicates and three technical replicates. A one-way ANOVA with Tukey’s multiple-comparison post-hoc test was used to compare dose effects within timepoints or a single dose across timepoints. A 2-sample t-test was used to compare GEDI ratios between two timepoints.

## Expected Results and Discussion

This protocol demonstrates how to use the Genetically Encoded Death Indicator (GEDI) biosensor to measure therapy-induced cell death in cancer cells. We selected SF8628, a H3K27M pediatric diffuse midline glioma (pDMG) cell line due to its documented intratumoral heterogeneity and prior use in radiosensitization and xenograft studies^10–15^. GEDI was stably transduced into SF8628 cells and FACS enriched for high mApple expression using appropriate positive and negative controls (**Supplemental File S3**.

To establish an empirical GEDI threshold for cell death, we first exposed cells to a high radiation dose (25 Gy), selected to reliably induce cell death and ensure a strong GEDI signal based on previous studies^16–18^. Cells were imaged longitudinally at defined intervals post-irradiation (**Figure 2A**). In these images, cells undergoing cell death displayed bright GC150 activation (yellow), while others remained GC150-negative (red only). To validate that the visual fluorescence shifts corresponded to quantitative shifts in the GEDI ratio, we tracked individual cells from the same field of view across the time course and quantified their GC150/mApple intensities using ImageJ (**Figure 2B**). Radiosensitive cells showed a rapid rise in GEDI ratio as they underwent cell death, whereas surviving cells maintained stable values. For population-level analysis, mApple-based object masks were used to quantify the fluorescence intensity of the mApple and GC150 channels for all cells in each field of view, using the ImageJ script provided in the protocol (**S1**). At 25 Gy, GEDI ratios were significantly higher at 24 hours post-irradition (hpi) compared to baseline (**Figure 2C**). Finally, a previously published method was applied to calculate a GEDI death threshold by comparing mean GEDI ratios of cells at 0 hpi (live) and 24 hpi (dead), yielding a threshold of 1.02 (**Figure 2D**)^9^. This empirically defined threshold separated live from dead cells and was used to classify cell death in subsequent radiation-based experiments with SF8628 cells. Together, these results demonstrate that radiation-induced stress can be quantified with the GEDI ratio in H3K27M-pDMG cells and provide a framework for assessing the heterogeneity of radiation response at clinically relevant doses.

**Figure 2:**
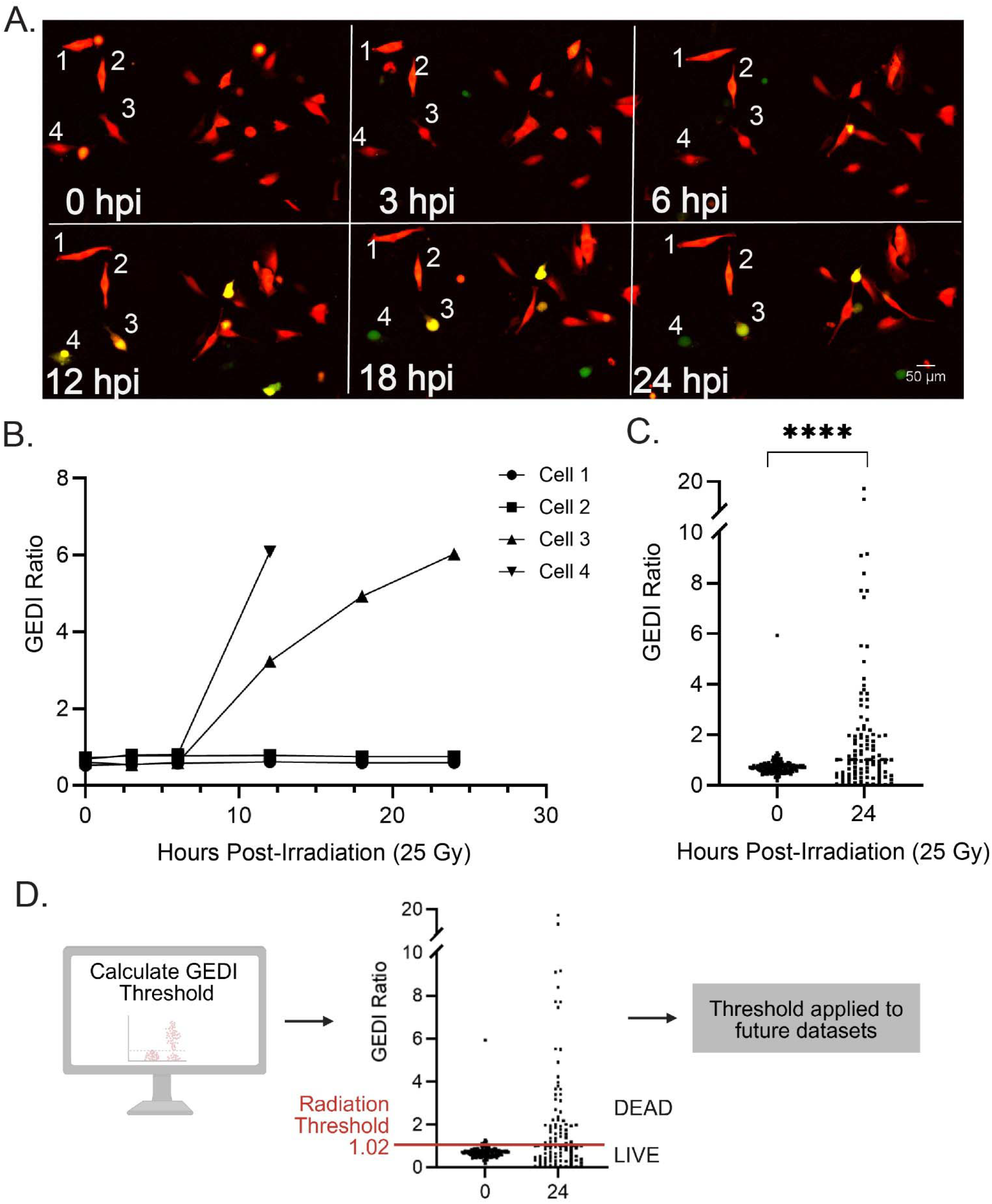
Determination of a GEDI ratio threshold for radiation-induced cell death. (**A**) Representative time-lapse montage of SF8628 GEDI-expressing cells exposed to 25 Gy x-ray irradiation and imaged at 0, 3, 6, 12, 18, and 24 hours post irradiation (hpi). Numbered cells (1-4) were tracked in B. (**B**) GEDI ratio traces of individual cells from (A) quantified at the indicated timepoints post irradiation. (**C**) GEDI ratio of cells in a 10 x 10, 20X montage field of view at 0 hpi (n = 309) and 24 hpi (266 cells).^****^p<0.0001, unpaired t-test. (**D**) The GEDI ratio threshold for cell death was calculated from (C) using *GEDI Threshold = [(Mean GEDI Dead – Mean GEDI Live) x 0*.*25] + [Mean GEDI Live]* and found to be 1.02. This calculated GEDI threshold is used to determine live and dead cell populations in future experimental conditions. Linear adjustments in ImageJ were made for representative figure images to highlight cellular features of interest. All quantification was done on raw images.

Having established the GEDI threshold for radiation-induced cell death, we next compared the sensitivity of GEDI with a commonly used endpoint assay for apoptosis, Caspase 3/7 Glo^19^. SF8628 GEDI-expressing cells were treated with 8 Gy irradiation, and both GEDI time-lapse imaging and caspase 3/7 signal were measured in parallel from 0 to 36 hpi. Cells were classified as dead if their GEDI ratio exceeded the threshold. (**Figure 3A**). GEDI analysis revealed a clear, time-dependent increase in the number of dying cells, with a strong linear relationship between the number of GEDI-dead cells and hours post-irradiation (R^2^ = 0.95) (**Figure 3B**). In contrast, caspase signal remained low until 12 hpi and only showed a significant increase at 36 hpi compared to earlier timepoints (**Figure 3C**). Although caspase activity and GEDI readouts were significantly correlated with radiation-induced cell death, caspase activity showed a greater variability (R^2^ = 0.71), reflecting its delayed and less consistent detection of death events. These results demonstrate that GEDI captures radiation induced cell death with higher temporal resolution than caspase-based assays and support its use as a complementary tool alongside other sensitive cell death indicators such as Annexin V.

**Figure 3:**
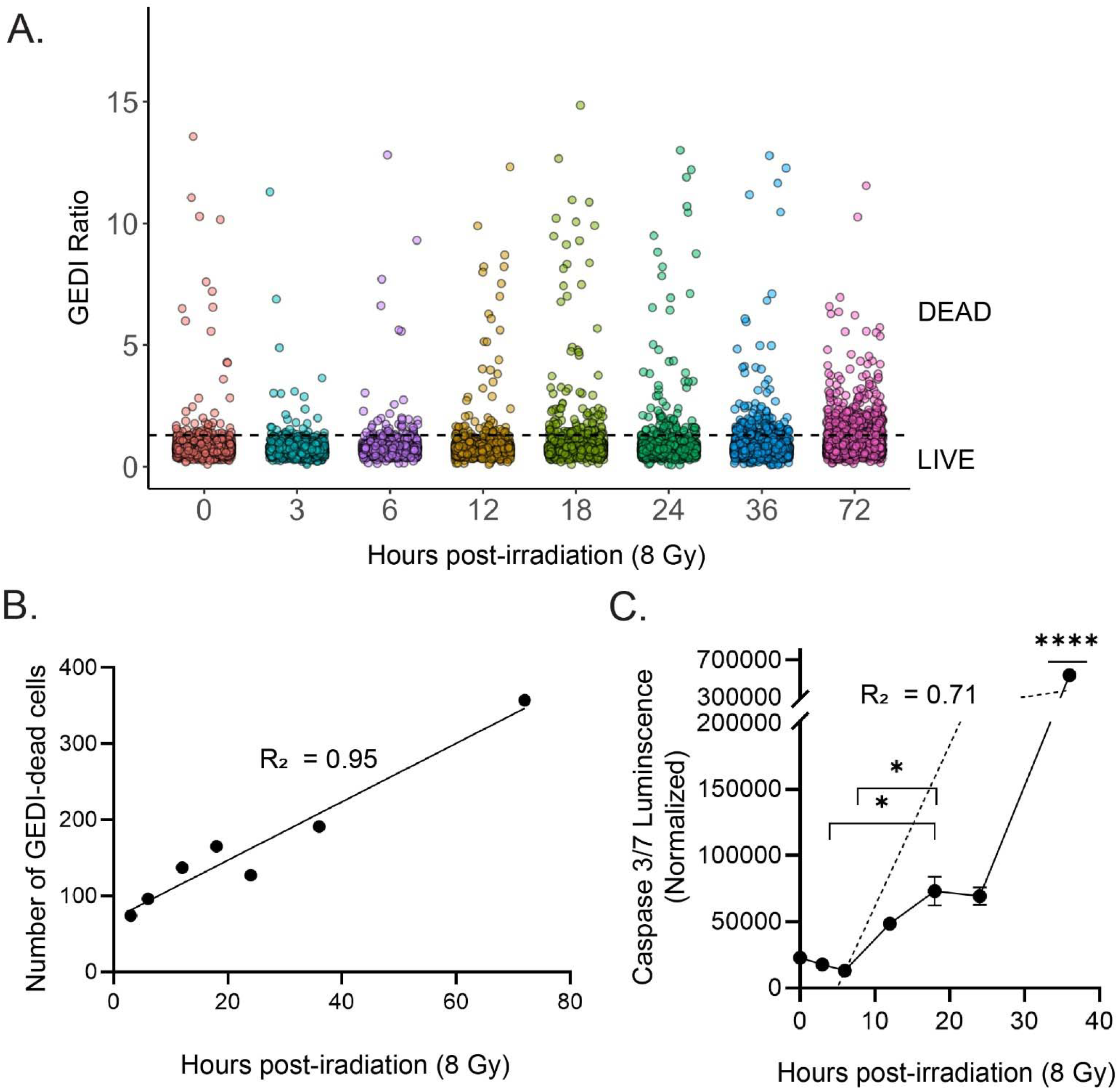
GEDI captures radiation-induced cell death with higher temporal resolution than conventional caspase 3/7 detection. (**A**) GEDI ratios of SF8628 cells at 0-72 h after a radiation dose of 8Gy. Each dot represents a single cell (n = 1600-2500 cells per timepoint). The dotted line indicates the radiation-induced GEDI threshold for death determined in Figure 2. (**B**) Linear regression of the number of GEDI-dead cells (above the threshold in A). Pearson correlation was used, R^2^ = 0.944, p < 0.0001. (**C**) Caspase 3/7 Glo luminescence at similar timepoints. Signal was normalized to media-only wells. The dashed line represents a linear regression fit (R^2^ = 0.71). Error bars are SEM (n = 3). Significance was determined by one-way ANOVA followed by Tukey’s post-hoc test, comparing all timepoints. Brackets indicate specific pair-wise comparisons that reached significance (^*^p < 0.05), while ^****^ indicates a timepoint significantly different from all others (p < 0.0001).

Chemotherapy remains a standard treatment strategy for many cancers, but heterogenous tumor cell populations often contain resistant subclones that survive therapy and drive relapse^20^. To establish a drug-induced GEDI threshold in parallel with our radiation model, we used JQ1, an inhibitor of the bromodomain and extraterminal domain (BET) family of proteins^21^. BET inhibition has shown efficacy in pre-clinical models of H3K27M-pDMG, yet recent studies of the subclonal diversity in these tumors highlight the need to distinguish drug-sensitive from drug-resistant cells at the single-cell level^12,22^. As an initial step, we performed two conventional endpoint assays to identify an appropriate lethal dose for threshold calibration. Caspase 3/7 Glo showed a significant increase in cell death and Cell Titer Glo revealed a significant decrease in cell viability at 50 µM JQ1 (**Supplemental Figure 4**). We then treated SF8628 GEDI expressing cells with 50 µM JQ1 and observed a significant increase in the GEDI ratios between 12-36 hours post-treatment (hpt) compared to baseline (**Figure 4A**). The highest mean GEDI ratio from this dataset was used to define the JQ1-induced death threshold (1.06). This threshold was slightly higher than the radiation-induced threshold (1.02), and peaked later (36 h vs 18 h), highlighting the need for treatment-specific calibration. We next evaluated lower, screening-relevant doses of JQ1 (1 µM and 10 µM). GEDI detected dose-dependent increases in death as early as 24 hpt (**Figure 4B**), and linear regression analysis confirmed strong temporal correlations at both doses (1 µM; R^2^ = 0.958, 10 µM; R^2^ = 0.993) (**Figure 4C**). Importantly, resistant cells that failed to cross the GEDI threshold were identified as early as 24 hpt (**Figure 4D**). These results demonstrate the utility of GEDI in detecting drug-sensitive from drug-resistant cells in bulk populations at the single-cell level, enabling rapid and quantitative screening of therapeutic responses.

**Figure 4:**
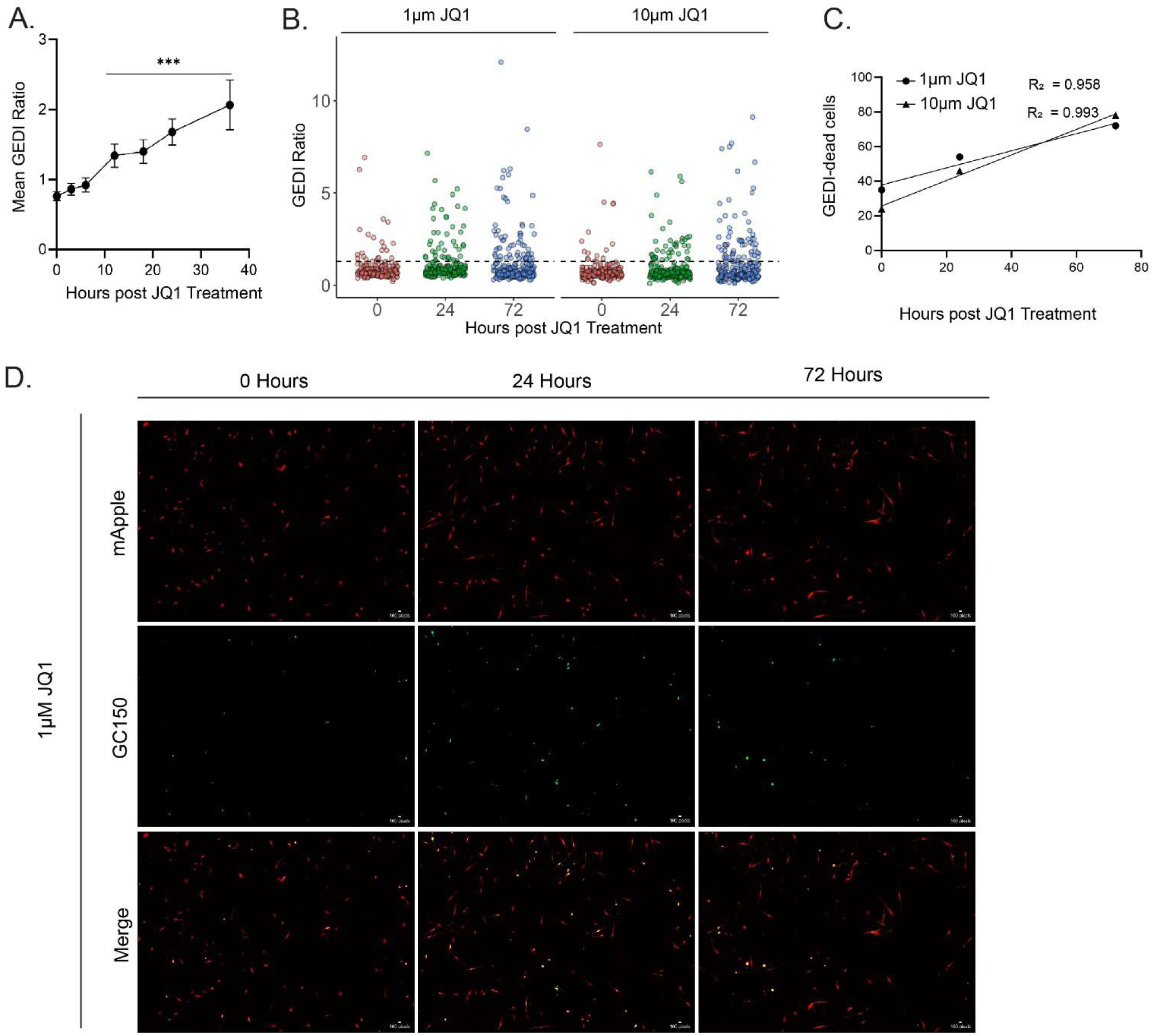
GEDI detects drug-induced cell death and resistance in SF8628 glioma cells. (**A**) Mean GEDI ratios of SF8628 cells treated with 50 µM JQ1 over 0-36 h. The peak ratio (1.06) was used to define the JQ1-induced GEDI threshold. Error bars are 95% confidence intervals, n = 419 cells. Significance by one-way ANOVA followed by a Tukey’s post-hoc; each timepoint compared to 0 h. ^***^p < 0.001 (**B**) Dot plot of GEDI ratios from SF8628 cells treated with 1 µM and 10 µM JQ1 at three timepoints. Dashed line represents GEDI threshold from (A). (**C**) Linear regression of GEDI-dead cells over time for 1 µM (R^2^ = 0.958) and 10 µM (R^2^ = 0.993) JQ1 treatments. (**D**) Representative images of SF8628 GEDI cells treated with 1uM JQ1 at 0, 24, and 72 hours post treatment. Linear adjustments in ImageJ were made for representative figure images to highlight cellular features of interest. All quantification was done on raw images.

High-throughput imaging generates large datasets that are difficult to analyze manually, making automated cell tracking essential for extracting dynamic information at the single-cell level. Advances in tracking software have greatly expanded the ability to study heterogeneous cell behaviors over time^23,24^. To test whether GEDI could be incorporated into such workflows, we applied TrackMate, a user-friendly automated cell tracking plugin available on ImageJ^25^. Integrating GEDI with automated tracking allowed us to generate high-confidence single-cell survival analyses that capture both intrinsic heterogeneity in death timing and potential extrinsic influences from neighboring cells. Our workflow, (**Figure 5A**; Part IV of the io.protocol) consisted of importing hyperstacked images into TrackMate and exporting GEDI ratio data for downstream analysis. Using this pipeline, we quantified GEDI live and dead cells across radiation doses of 2, 8, and 25 Gy over multiple time points (**Figure 5B**). Representative cell tracks from the 8 Gy dataset were also inspected manually to confirm tracking accuracy (**Figure 5C**). These results demonstrate that GEDI is compatible with open-source automated cell tracking platforms, enabling scalable, unbiased analysis of therapy-induced cell death dynamics.

**Figure 5:**
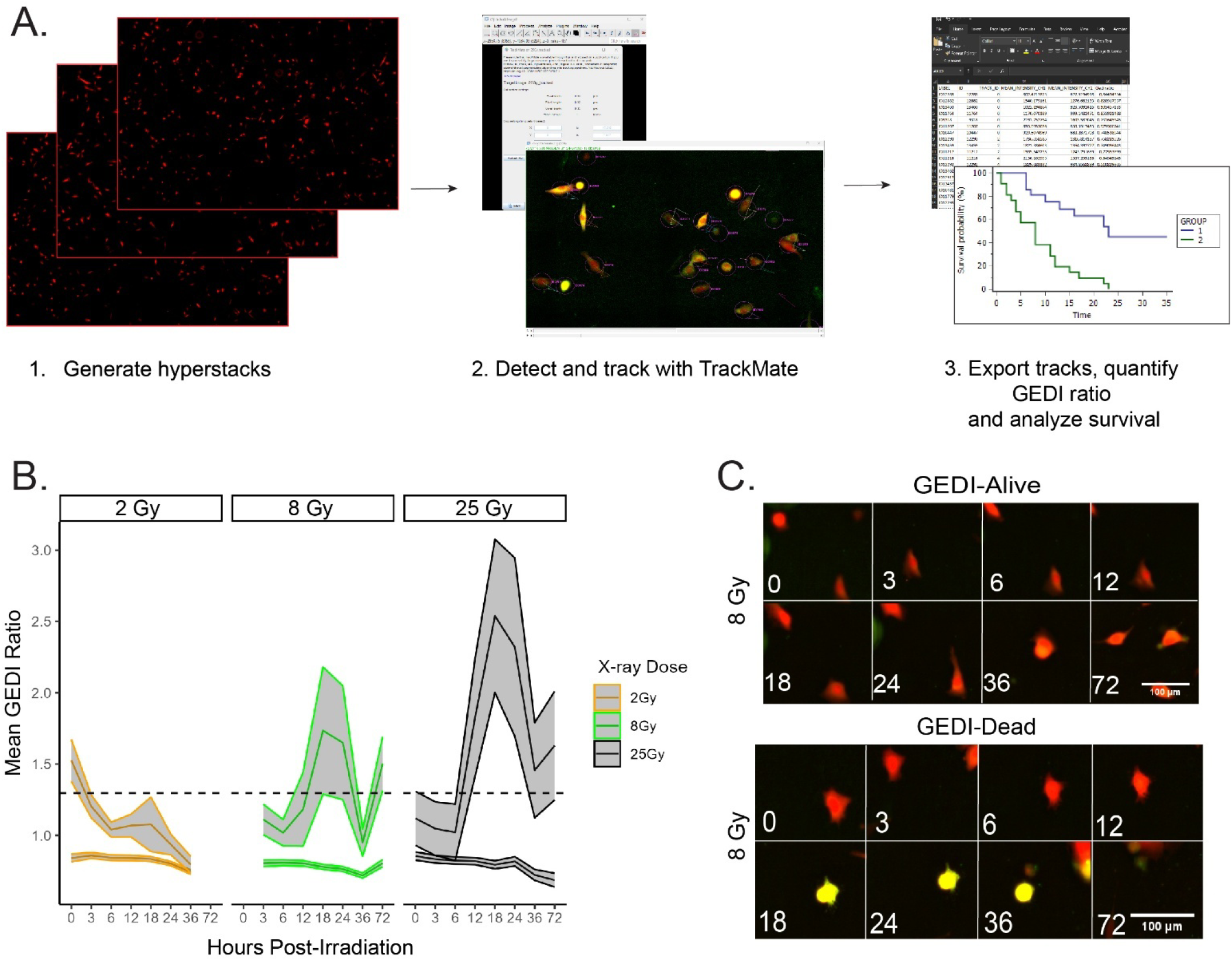
GEDI is compatible with Trackmate for automated quantification and live cell tracking. **(A)** Workflow of single-cell tracking of SF8628 GEDI-expressing cells using Trackmate. Hyperstacks are generated, cells are detected and tracked, and GEDI ratios are quantified to generate survival data. **(B)** Mean GEDI ratios of cells treated with 2, 8, or 25 Gy radiation and tracked using Trackmate for seven time intervals post-irradiation (n = 200-300 cells per condition). Shaded areas represent 95% confidence intervals. The dashed line represents the radiation-induced GEDI threshold (see Figure 2). **(C)** Representative Trackmate-generated montage images of single tracks from the 8 Gy irradiated cells. Shown are examples of a cell that remains below the GEDI threshold (“Alive”) and a cell that exceeded the threshold (“Dead”). Linear adjustments in ImageJ were made for representative figure images to highlight cellular features of interest. All quantification was done on raw images.

GEDI’s ability to classify live and dying cells extends beyond thresholding into functional and molecular applications (**Figure 6**). GEDI can be incorporated into high-throughput drug testing to quantify death kinetics, dose-response relationships, and drug or radiation synergies, providing rapid and quantitative readouts of therapeutic efficacy. It also allows for the identification and enrichment of treatment-resistant subclones, a crucial step in understanding tumor heterogeneity and how rare cells survive to drive relapse. These resistant populations can be tracked over time, rechallenged with secondary treatments, or directly compared to dying cell populations to uncover the mechanisms of persistence. Finally, GEDI-expressing cells can be advanced into downstream biological and molecular assays, including xenograft, explant, or organoid models to test resistance within a tumor microenvironment, or cells can be subjected to transcriptomic, epigenomic, or gene perturbation studies to define pathways of therapeutic response. By linking dynamic cell death measurements with functional and molecular characterization, GEDI provides a versatile platform to dissect therapeutic response, resistance, and clonal evolution in cancer.

**Figure 6:**
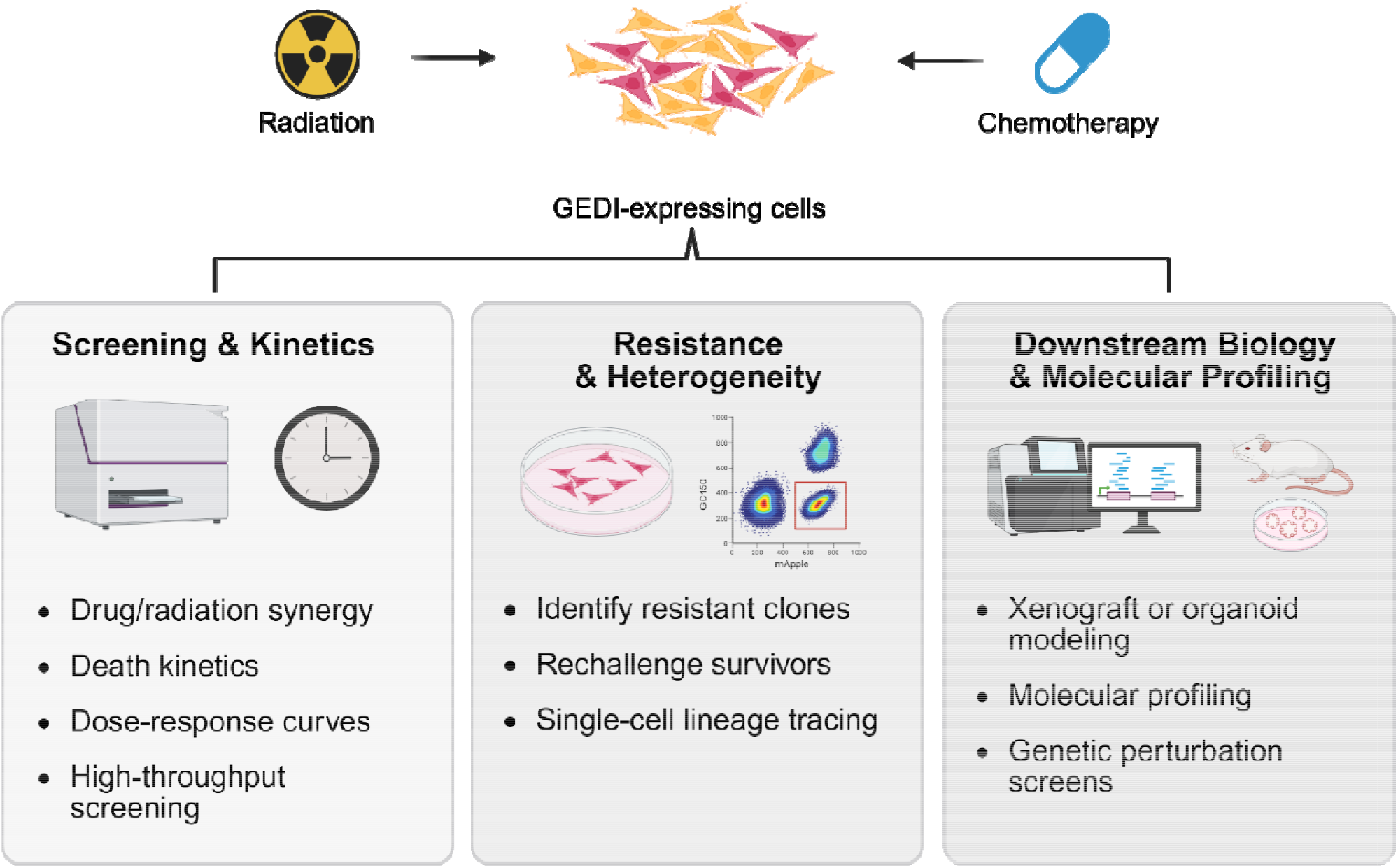
Applications of GEDI-expressing cells in cancer research. After treatment with radiation or chemotherapy, GEDI-expressing cells can be routed into downstream workflows to quantify therapeutic efficacy, dissect resistance mechanisms, and inform cancer biology and treatment design at single-cell resolution.

In summary, this study demonstrates that a biosensor originally developed to detect neurodegeneration can be repurposed to screen for radio- and chemo-resistant cancer cells. Using SF8628, a treatment-naive H3K27M-pediatric glioma cell line, we established radiation- and drug-induced thresholds for cell death and showed that GEDI provides both temporal and spatial resolution of death events in heterogenous populations. We further demonstrate that GEDI is compatible with open-source automated single-cell tracking, enabling high-throughput single-cell survival analyses.

While GEDI offers a highly sensitive approach for screening radiation- and chemotherapy-resistant cancer cells, there are several limitations to consider. Imaging parameters must be kept consistent from GEDI threshold determination through experimental runs to ensure accurate quantification. We have utilized Gen5’s image processing tools, however, ImageJ offers several similar algorithms. Careful calibration of exposure settings is required for optimal signal-to-noise; for example, best results were achieved in our system when GC150 exposure was set at two times the exposure of mApple. Imaging systems must allow repeated acquisition of the same field of view to support time-lapse imaging. Empirical determination of a GEDI threshold should also be done at each treatment condition, as shown by the slight differences between radiation- and drug-induced thresholds and the time required to reach a maximum. Additionally, combination treatments may need further calibration. Finally, TrackMate’s preset spot and tracking detectors can generate errors, so manual curation of cell tracks or the use of more advanced tracking algorithms are recommended to improve accuracy.

Despite these limitations, GEDI markedly advances the ability to detect therapy-induced cell death at single-cell resolution. Compared with traditional endpoint assays, GEDI captures early and dynamic death events, distinguishes resistant subclones, and provides a flexible platform that can be paired with downstream functional and molecular analyses. This sensitivity has the potential to accelerate preclinical drug and radiation screening and to deepen our understanding of heterogeneity and resistance mechanisms in cancer.

## Ethics declarations

This study does not use humans or animals.

## Supporting Information

All supporting information is included in supplemental files or on Github. Raw outputs from FIJI analysis, example images, and example FIJI and R scripts for GEDI quantification are provided on the Github repository “GEDI_Cancer”.

## Acknowledgments

The authors acknowledge the University of Kentucky the Flow Cytometry and Immune Monitoring core for the use of their facilities. We thank Steven Finkbeiner (The Gladstone Institutes) for the GEDI plasmid.

## Author’s contributions

**Viral Oza**: conceptualization (lead), data curation (lead), formal analysis (lead), investigation (lead), methodology (lead), project administration (lead), writing-original draft (lead), writing-review and editing (lead). **Collin Williams** data curation (supporting), methodology (supporting), writing-original draft (supporting), writing-review and editing (supporting). **Jessica Blackburn** conceptualization (lead), funding acquisition (lead), formal analysis (supporting), project administration (lead), writing-review and editing (lead).

