## Supplementary Data 2-4 for "A live cell biosensor protocol for high-resolution screening of therapy-resistant cancer cells"

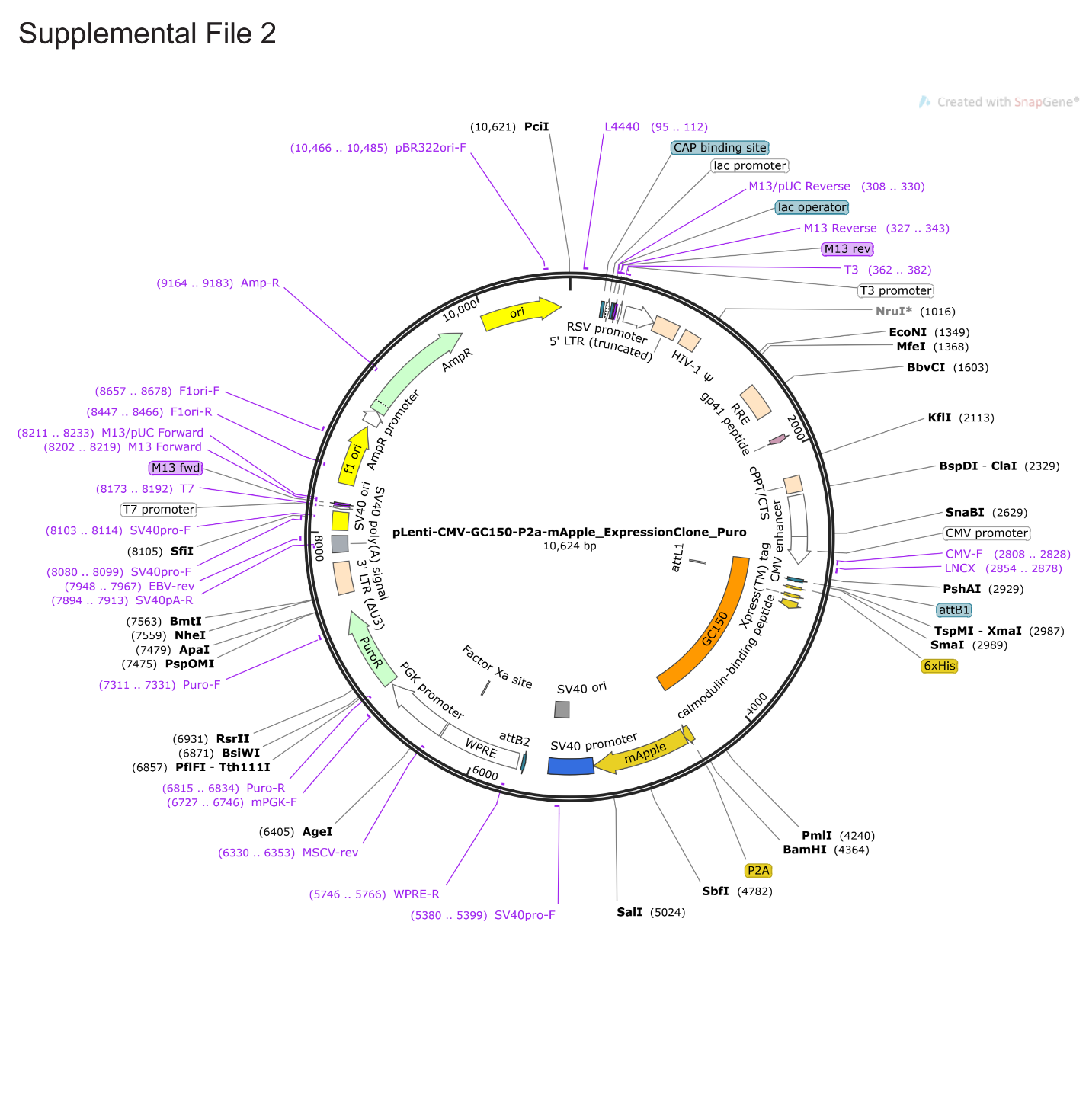

Supplemental File 2: Plasmid Map of GEDI construct.

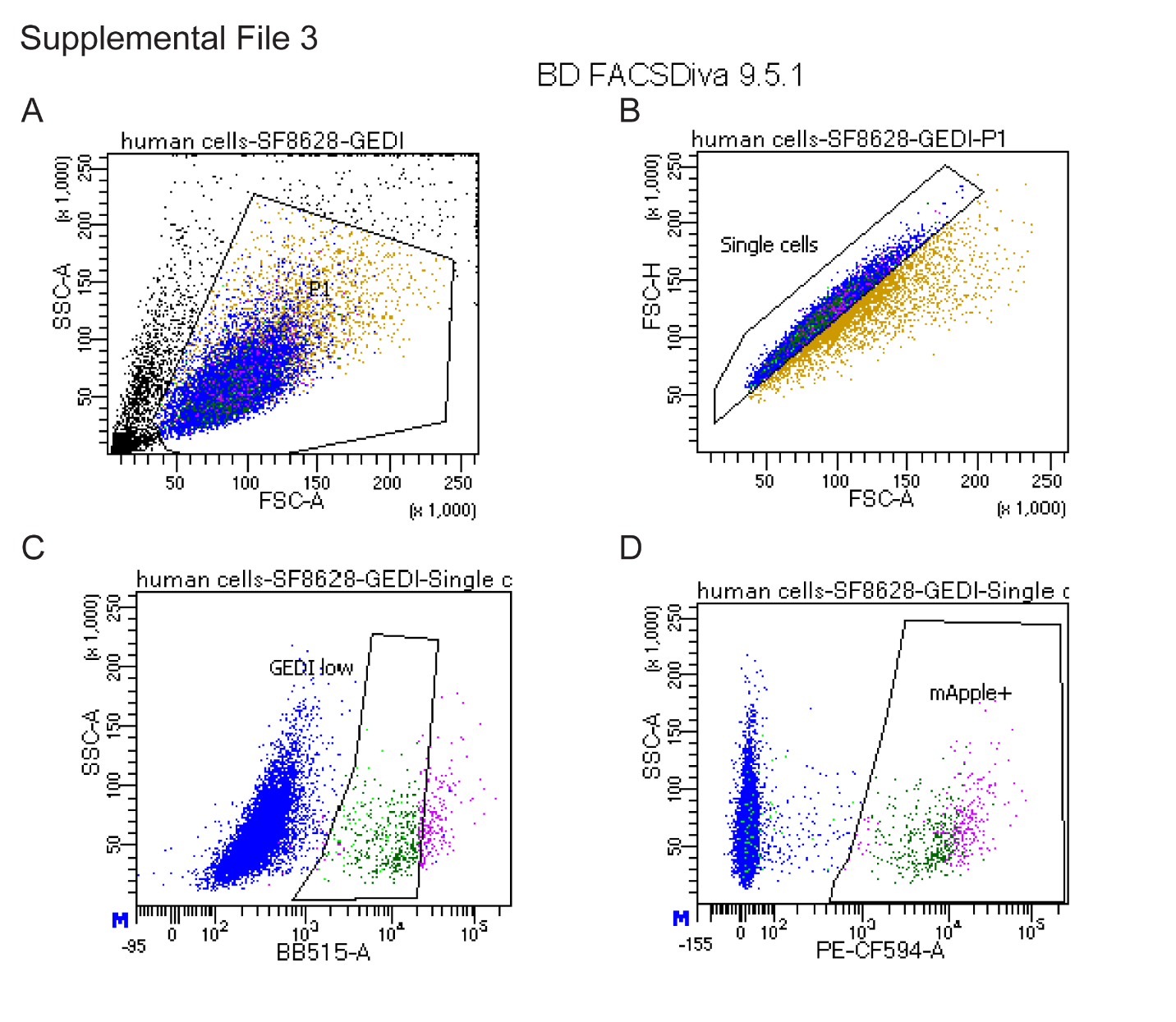

Supplemental File 3: FACS sorting of GEDI transduced SF8628 cells. (top left) Gating scheme of side scatter vs. forward scatter signal, (top right) Gating scheme of forward scatter height vs forward scatter width, (bottom left) Gating scheme capturing GC150 low expressing cells after gating out positive control (heat shocked cells), (bottom right) gating scheme of highly expressing mApple positive cells.

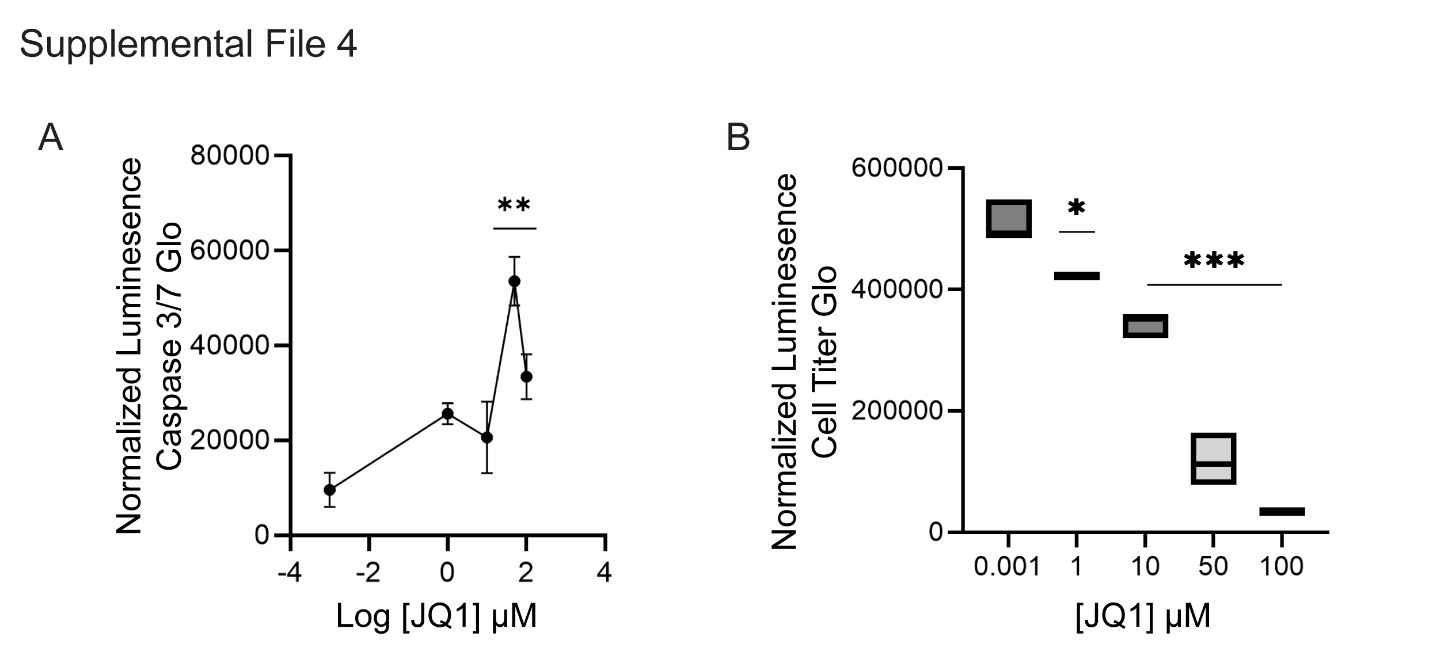

Supplemental File 4: Caspase 3/7 Glo and Cell Titer Glo inform selection for GEDI threshold calibration (A) Cell death data showing normalized caspase 3/7 signal from SF8628 cells treated at a range of JQ1 concentrations 24 hours post JQ1 treatment. Error bars represent SEM, n = 3. Significance as determined by a one-way ANOVA followed by a Tukey’s post hoc, ** p<0.01. (B) Cell viability data showing normalized Cell Titer Glo signal from SF8628 cells treated at a range of JQ1 concentrations 24 hours post JQ1 treatment. Error bars represent SEM, n = 3. Significance as determined by a one-way ANOVA followed by a Tukey’s post hoc,* p<0.05,***p<0.001
