## Supplemental Data 1 for "A live cell biosensor protocol for high-resolution screening of therapy-resistant cancer cells"

RESERVED DOI:
10.17504/protocols.io.eq2ly4d7qlx9/v1

Viral D. Oza[1,2], Colin S. WIlliams[1,2], Jessica S. Blackburn[1,2]

[1]Department of Molecular and Cellular Biochemistry, University of Kentucky, Lexington, KY, USA;
[2]Markey Cancer Center, University of Kentucky, Lexington, KY, USA

Viral Oza
University of Kentucky

**Protocol Info:** Viral D. Oza, Colin S. WIlliams, Jessica S. Blackburn . A live cell biosensor protocol for high-resolution screening of therapy-resistant cancer cells. **protocols.io** **https://protocols.io/view/a-live-cell-biosensor-protocol-for-high-resolution-g4xfbyxjp**

**Created:** July 07, 2025

**Last Modified:** September 15, 2025

**Protocol Integer ID:** 221895

**Keywords:** therapy resistance, cell death, biosensor, GEDI, genetically encoded

**Funders Acknowledgements:**
**National Institutes of Health**
Grant ID: R37CA227656
**National Institutes of Health**
Grant ID: F99CA294265
**Kentucky Pediatric Cancer Research Trust**
Grant ID: PON27282400002665
**National Institutes of Health**
Grant ID: P30CA177558

#### Abstract

The Genetically Encoded Death Indicator (GEDI) is a ratiometric, dual-fluorescence biosensor that enables real-time detection of cell death through calcium influx. Originally developed for use in neurodegeneration models, GEDI can be applied to cancer cells to quantify therapy-induced death at single-cell resolution. This protocol details how to generate GEDI-expressing cancer cell lines, empirically determine stress-induced GEDI thresholds using radiation or chemotherapeutic agents, and perform time-resolved imaging and image analysis to track cell fate. This workflow is optimized for high-throughput drug and radiation screening in heterogeneous populations and is especially useful for identifying chemo- and radio-resistant subclones. Key limitations include the need for empirical GEDI threshold calibration for each treatment condition and careful standardization of imaging parameters. The protocol outputs include GEDI ratio values, single-cell time-of-death annotations, and whole-cell morphological data in parallel, which can be linked to downstream applications such as FACS-based isolation of live or dying subpopulations, transcriptomic profiling of resistant clones, or in vivo validation using xenografts or organotypic slice culture.


#### Guidelines

General considerations:
For best signal to noise ratios set the GC150 exposure to 2X that of mApple exposure
Sparse seeding of cells will enable greater accuracy of tracking algorithms
Name Image files with relevant metadata for straightforward dataframe manipulation downstream

### Materials

Biological Materials
1. H3K27M-pDMG cell line SF8628 (SCC127, Millipore Sigma)
2. HEK293T celll line (12022001, Sigma)

Reagents
1. 0.25% Trypsin (25300062, Life Technologies)
2. DMEM 1X with 4.5g/L Glucose (D6546, Sigma), L-Glutamine (07100, Stem Cell Technologies), & Sodium Pyruvate (07000, Stem Cell Technologies)
3. FBS (S11150H, R&D Systems)
4. PBS 1X (PBD500-CS, Sigma)
5. Lipofectamine (L30000015, Thermo Scientific)
6. Lenti-X (631231, Takara Bio)
7. HSBSS (55037C, Sigma)
8. Trypan Blue (C10283, Thermo Scientific)
9. pLenti-CMV-Puro destination vector (17452, Addgene)
10. psPAX2 (12260, Addgene)
11. pMD2.g (12259, Addgene)
12. Invitrogen Gateway technology (11791020, Life Technologies)
13. TransIT reagent (2300, Mirus Bio)
14. Lenti-X (631231, Takara Bio)
15. Caspase-Glo 3/7 Reagent (PAG8091, VWR)
16. Cell Titer-Glo Luminescent Cell Viability Assay Reagent (PAG7572, VWR)
17. Optimem (31985070, Fisher Scientific)

Cell culture media
1. DMEM 1X with 4.5g/L Glucose
2. 10% FBS
3. 1X L-Glutamine
4. 1X Sodium Pyruvate

Laboratory Supplies and equipment
1. Biotek Lionheart
2. BD FACS Aria
3. 100mm tissue culture dish
4. Hemocytometer cell counting chamber
5. Computer with at least 7.9Gb memory
6. FACS tubes (60819-295, VWR)
7. FACS strainer (23A00A612, Thomas Scientific)
8. 15mL conicals
9. 96 well black wall plates (29444, VWR)
10. Biotek Synergy LX Multi-Mode Reader
11. Luminescent Filter Cube (1505003, Biotek)
12. 96 well white walled plates (29444, VWR)
13. LookOut Mycoplasma PCR Detection Kit (MP0035, Sigma)

#### Protocol materials

- pLenti CMV Puro DEST **addgene Catalog** #17452
- Gateway LR Clonase II Enzyme mix **Thermo Fisher Catalog** #11791020
- EcoR1-HF **NEB Catalog** #R3101S
- Caspase 3/7 Glo **VWR International (Avantor) Catalog** #PAG8091
- 0.25% Trypsin **Life Technologies Catalog** #25300062
- Lenti-X **Takara Bio Inc. Catalog** #631231
- psPAX2 **addgene Catalog** #12260
- pMD2.g **addgene Catalog** #12259
- TransIT **Mirus Bio Catalog** #2300
- Optimum **Fisher Scientific Catalog** #31985070
- 1x PBS (Phosphate Buffered Saline) **Merck MilliporeSigma (Sigma-Aldrich) Catalog** #PBD500-CS
- HSBSS **Merck MilliporeSigma (Sigma-Aldrich) Catalog** #55037C
- Cell Titer Glo **VWR International (Avantor) Catalog** #29444
- FBS **R&D Systems Catalog** #S11150H
- 1X L-Glutamine **STEMCELL Technologies Inc. Catalog** #07100
- 1X Sodium Pryvate **STEMCELL Technologies Inc. Catalog** #07000
- HEK 293T **ATCC Catalog** #ATCC CRL-3216)
- DMEM 1X w 4.5g/L Glucose **Merck MilliporeSigma (Sigma-Aldrich) Catalog** #D6546

#### Safety warnings

Ensure proper handling when using lentivirus and lentiviral infected materials.

#### Ethics statement

This study does not use humans or animals.

#### Part I: Prepare GEDI lentivirus, transduce and sort cells

1w 6d 0h 50m

1 To generate GEDI lentivirus, the pMe:GC150-p2A-mApple construct (gift from the Finkbeiner Lab, Gladstone Institutes) was cloned into a pLenti CMV Puro DEST **addgene Catalog** #17452 vector using Gateway LR Clonase II Enzyme mix **Thermo Fisher Catalog** #11791020, according to manufacturer's instructions. Clones were selected based off an EcoR1-HF **NEB Catalog** #R3101S restriction digest and confirmed through whole plasmid sequencing. 3d

2 Start culture of HEK 293T **ATCC Catalog** #ATCC CRL-3216) cells in DMEM 1X w 4.5g/L Glucose **Merck MilliporeSigma (Sigma-Aldrich) Catalog** #D6546 with 10% FBS **R&D Systems Catalog** #S11150H , 1X L-Glutamine **STEMCELL Technologies Inc. Catalog** #07100 , and 1X Sodium Pryvate **STEMCELL Technologies Inc. Catalog** #07000 .

3 Lentivirus was produced by transfecting $4 * 10^6$ HEK 293T cells in a 10cm plate with the following three plasmids, pSPAX2 **addgene Catalog** #12260 , pMD2.g **addgene Catalog** #12259 , and GEDI expression plasmid in the following ratio $1 : 0.1 : 0.9$ in 250 µL of Optimem. Mix 12 µL of TransIT **Mirus Bio Catalog** #2300 with 238 µL of Optimum **Fisher Scientific Catalog** #31985070. Incubate both mixtures for 00:05:00 and room temperature. 5m

3.1 Mix the two solutions together and pipette up and down. Incubate 00:20:00 at room temperature. Add total mixture drop wise to 10cm plate. Change media next day. 20m

3.2 Viral supernatant was collected at 48:00:00 and 72:00:00 post-transfection. Viral supernatant was spun 00:05:00 at 300 x g. Supernatant was collected and filtered through 0.45 micron filter. Viral supernatant concentrated using Lenti-X **Takara Bio Inc. Catalog** #631231 Concentrator and stored in 10 µL aliquots at -80 °C . 2d 0h 5m

4 Seed cells at 10000 cells per well and test a range of lentivirus doses ranging from 0.25 µL - 2 µL per well.

5 Check for expression and toxicity using brightfield and a filter set amenable to mApple excitation/emission. We used a Lionheart FX imager with the following LEDs and filter cube sets:

| Equipment | |
|---|---|
| **RFP filter cube** | NAME |
| filter cube | TYPE |
| Biotek | BRAND |
| PN:1225103 | SKU |

| Equipment | |
|---|---|
| **GFP filter cube** | NAME |
| filter cube | TYPE |
| Biotek | BRAND |
| PN:225101 | SKU |

| Equipment | |
|---|---|
| **523nm LED cube** | NAME |
| LED cube | TYPE |
| Biotek | BRAND |
| PN:1225003 | SKU |

| Equipment | |
|---|---|
| 465nm LED cube | NAME |
| LED cube | TYPE |
| Biotek | BRAND |
| PN: 1225001 | SKU |

| Equipment | |
|---|---|
| Lionheart FX Imager | NAME |
| Microscope | TYPE |
| Biotek | BRAND |

6 After lentivirus dose has been determined, transduce cells in a 10cm dish at ~ 50 % confluency

7 After 120:00:00 , start feeding cells and prepare test cells for sorting using a 5d

| Equipment | |
|---|---|
| iCyt-Sony Cell Sorter | NAME |
| cell sorter | BRAND |

8 For cells to be sorted, repeat the following steps (Part 2: 15-18).

9 During resuspension, resuspend cells in HSBSS **Merck MilliporeSigma (Sigma-Aldrich) Catalog #55037C** warmed to

37 °C

10 Dilute cells to $1 * 10^6$ cells/mL and strain through a FACS strainer into a FACS tube and place on ice.

11 Set up a negative control cell population from non-transduced cells. Set up a positive control by resuspending $5.0 * 10^5$- $1.0 * 10^6$ GEDI transduced cells in 1 mL and heat shock at 42 °C for 00:10:00 , then strain through FACS strainer into FACS tube and place on ice. 10m

12 Optional: For a secondary positive control spike in NaN3 for a total concentration of [M] 2 % (v/v) for 00:10:00 before sorting begins. 10m

13 During sorting, gate for cells with high mApple expression and low GC150 expression. Set a negative gate on unstained cells (low mApple, low GC150) and a positive gate on dead cells (high mApple, high GC150).

14 Let cells recover for several days after sorting, at least 72:00:00 before using downstream experiments. 3d

#### Part II: Determine a lethal dose of drug of interest or x-ray radiation using Caspase 3/7 Glo or Cell Titer Glo

10m

15

When cells are ~ 70 % confluency in a 10cm dish wash with

1x PBS (Phosphate Buffered Saline) **Merck MilliporeSigma (Sigma-Aldrich) Catalog** #PBD500-CS

, then aspirate PBS.

16 Add 2 mL of 0.25% Trypsin **Life Technologies Catalog** #25300062 directly to plate and incubate in cell culture incubator for 00:05:00 5m

17 Quench Trypsin with 4 mL of freshly warmed media and transfer to 15mL conical tube.

18 Spin 15mL conical tube for 00:05:00 at 300 x g and resuspend cell pellet in 1 mL of fresh media. 5m

19 Seed cells at a concentration adequate for downstream experiments.

#### Drug Treatment 4h

20 Seed cells at a concentration of lower 5000 cells per well for drug treatment using steps from Part 2: 15-18. Total plating volume will be 50 µL per well.

21 Wait at least 04:00:00 after plating or until cells have attached back to the plate. 4h

22 Add 50 µL of freshly prepared 2X drug directly to the well and pipette up/down 3X

23 Proceed to Caspase 3/7 or Cell Titer Glo step.

#### Radiation Treatment 4h

24 Seed cells at a concentration of lower 5000 cells per well for radiation treatment using steps from Part 2: 14-18. Total plating volume will be 100 µL per well.

25 Wait at least 04:00:00 after plating or until cells have attached back to the plate. 4h

26 Bring 2 plates to the irradiator in Styrofoam housing.

27 Place plate getting doses > 0Gy into irradiator and adjust settings to achieve 8Gy exposure.

27.1 On a XRAD 225XL beam parameters are as follows, 13.3mA, SSD = 40, 225kV. For 8Gy, set the dose to 718.6.

**Equipment**

| | |
|---|---|
| XRAD 225XL | NAME |
| Precision X-Ray | BRAND |

28 Proceed to Caspase 3/7 or Cell Titer Glo step.

#### Cell Titer Glo

12m

29 Take tissue culture plate out and cool to room temperature before adding reagent.

30 At appropriate timepoint prepare Cell Titer Glo **VWR International (Avantor) Catalog** #29444 (CTG) reagent at a $1:3$ dilution in sterile 1X PBS in a total volume enough for 100 µL per well.

31 With the 100 µL of media in the well, add 100 µL of CTG to each well, pipette up/down 3X and shake, covered in foil, on orbital shaker for 00:10:00

10m

32 Let stand for 00:02:00 at room temperature and then read in plate reader with Luminescence detection mode.

2m

| Equipment | |
|---|---|
| Biotek Synergy LX Multi-Mode Reader | NAME |
| Biotek | BRAND |

| Equipment | |
|---|---|
| Luminescent Filter Cube | NAME |
| Biotek | BRAND |
| 1505003 | SKU |

#### Caspase 3/7 Glo

30m

33 At appropriate timepoint prepare Caspase 3/7 Glo **VWR International (Avantor) Catalog** #PAG8091 reagent in a total volume for 100 µL per well.

34 With the 100 µL of media in the well, add 100 µL of Caspase 3/7 Glo reagent to each well and pipette up/down 3X. Cover and incubate at 37 °C for at least 00:30:00 , can go up to 2 hours if necessary. 30m

35 Read in plate reader with Luminescence detection mode.

#### Part III: Determine GEDI threshold for cell death

36 Seed cells at a concentration of lower 2500 cells per well or lower to allow adequate cell tracking using steps from Part 2: 15-18.

37 Image GEDI expressing cells before stress stimulus.

37.1 Imaging of GEDI is done using filter sets described above, Part 1: 4.

38 Use filter sets described in Part 1: 5 for imaging. Exposure times will vary, recommended exposure settings are 150ms for mApple and 300ms for GC150 with a low-medium Gain setting.

39 Setup a Gen5 protocols recommended with the following considerations:

39.1 Determine optimal mApple exposure without saturating pixels.

39.2 Set GC150 exposure to twice that of mApple.

39.3 Set magnification to 20X for best signal to noise ratio.

39.4 Capture 10 × 10 montage of each well.

39.5 Use the following data reduction steps for post image processing. Background Subtraction (Rolling point-point spread function), Deconvolution (Gaussian), Stitching (linear blend without downsizing)

40 Induce Radiation and/or Drug Stress to cells.

40.1 Add appropriate amount of 2X drug in a 100uL volume and add to 100uL of volume in the plate.

40.2 If irradiating, transport plate to irradiator in Styrofoam housing, expose to appropriate amount of x-ray and return to cell culture incubator or the microscope.

41 Continue imaging GEDI expressing cells at timed intervals after stress induction.

#### Image J Data Analysis

42 Quantify the GEDI ratio per object (cell) in each image by running a script (attached) with these basic principles:

43 1) Threshold objects based on the mApple intensity. (FIJI script)

44 2) Use these object masks to measure the fluorescence intensity of the mApple and GC150 signal. (FIJI script)

44.1 FIJI Script:

```
//This FIJI script is designed to take RFP and GFP images from
GEDI data and output 1 CSV
//quantifying the fluorescent intensities from both channels

//It is specifically written for images of SF8628 cells that have
been exposed to 25 Gy and seeded //in 96 well plates at a 2500
cell per well density

//Designate where/what directory/folder the CSV output will be
saved
#@ File(label = "Output directory", style = "directory") output

//Make a list of images so it can be called by an integer value
list = getList("image.titles");


//The list integers start from 0, so double check which images are
opening first by showing the //array, this will dictate what image
in the list you are using below
Array.show(list);

//Select the open image window by its integer value from the list
to do further processing on, in //this case we are generating cell
masks from the RFP channel which is the second image in our
//image list, remember the list starts with 0 and the first image
in the order that we opened it is
// GFP, so to select the 2nd image that is open we would select
the list integer of "1" to designate selecting the RFP image.

selectWindow(list[1]);


//In this case we first want to duplicate the image to threshold
and leave the original untouched //because we will be overlaying
the mask
//on top of it to extract fluorescence intensity data

//Command to duplicate the open image
run("Duplicate...", " ");

//Manually set the threshold, the minimum value is one that should
be tested/iterated for what //works best to capture your cell mask
of interest
setThreshold(1139, 65535);

setOption("BlackBackground", true);
```

```
run("Convert to Mask");

///Analyze all the particles in your image to generate ROIs, we
will use these ROIs to place on the //original image and measure
the fluorescent intensity
//of the desired channel

run("Analyze Particles...", "size=150-Infinity display clear
include summarize add");
        //The parameters under analyze particles can be changed to
exclude a certain size, this will //need to be tested depending on
the size of your object and the magnification used,
//and what you are interested in. In our test case, we have
measured our cells and want the //minimum cutoff for particle
counting to be 150.
//If there are artifacts in your image that you can avoid using,
the size or feret diameter filter this is //a good place to
implement that

//You need to set the types of measurements that will be extracted
after measuring the image //mask, these can be changed, the best
way is to record a macros
//and select specifically what you want measured and then paste
that macros line here

run("Set Measurements...", "area mean standard min feret's
integrated area_fraction display redirect=None decimal=3");

//Clear the results becasue we do not want mask measurements in
the final CSV file
run("Clear Results");

//Select the original image, in this case it is the RFP image
selectWindow(list[1]);

//Overlay the ROIs from the ROI manager onto the image of interest
run("From ROI Manager");



//Now measure all of our ROIs on our new image
roiManager("Measure");

//Now we select the GFP image, in this case its the first image
that we opened, so the list index is //"0"
selectWindow(list[0]);
```

```
//Again, overlay the ROIs onto the selected GFP image "0" and
measure them
run("From ROI Manager");
roiManager("Measure");

//We want the title of the image in our filename, in this case the
title of the image has the expt //details that we can parse out
later with an R script
//so we use the following command to save the title of the image
into a variable called fileName that we can use to name our CSV of
the results file

fileName = getTitle();
  saveAs("Results", output + fileName + ".csv");

//Close all image windows
close("*")

//Clear results
run("Clear Results");
```

45 3) Divide the GC150 signal by the mApple signal to get the GEDI ratio. (EXCEL or R script)

45.1 Use the following equation to determine the GEDI ratio threshold for cell death where "mean GEDI ratio dead" is the mean GEDI ratio from cells that have been given a stress stimulus according to Part 1: step 10 and mean GEDI ratio live is the GEDI ratio of the aforementioned cells before the stress stimulus:
GEDI ratio threshold = [(mean GEDI ratio dead) – (mean GEDI ratio live)] ∗ 0.25 + [mean GEDI ratio live]

46 4) Bind CSVs into one file and by following the attached R Markdown File. (R script)

BIND_FIJI_CSVs.html 615KB

#### Part IV: Cell tracking with TrackMate

47 Open TrackMate. Must have hyperstack open. Screenshots of steps found below.

48 Use the loG detector for spot detection adjusting the spot threshold accordingly.

49 Use the LAP tracker to generate tracks.

50 Screenshots of steps used for TrackMate.

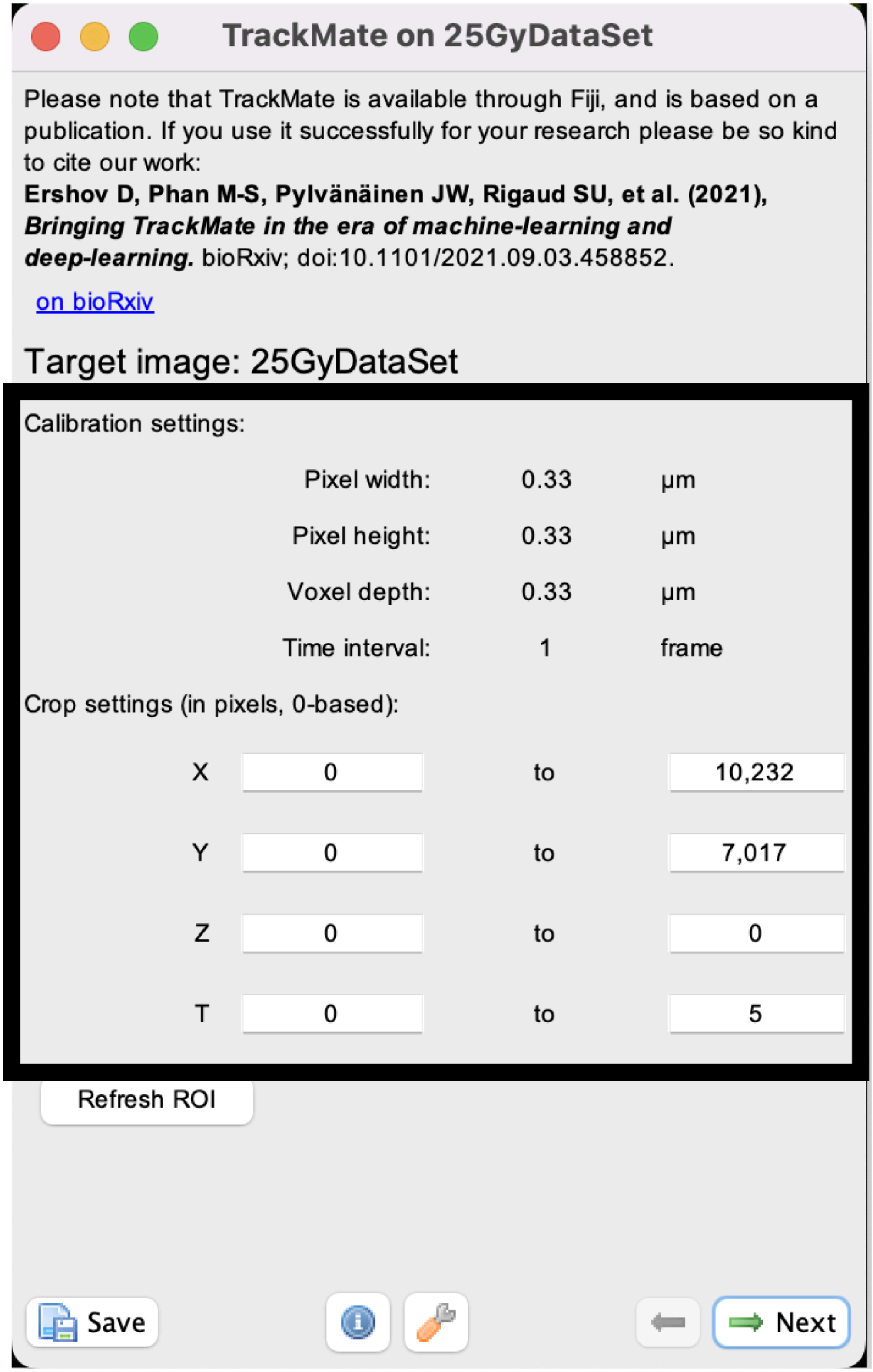


Check calibration settings and crop settings for user's liking. Click next.

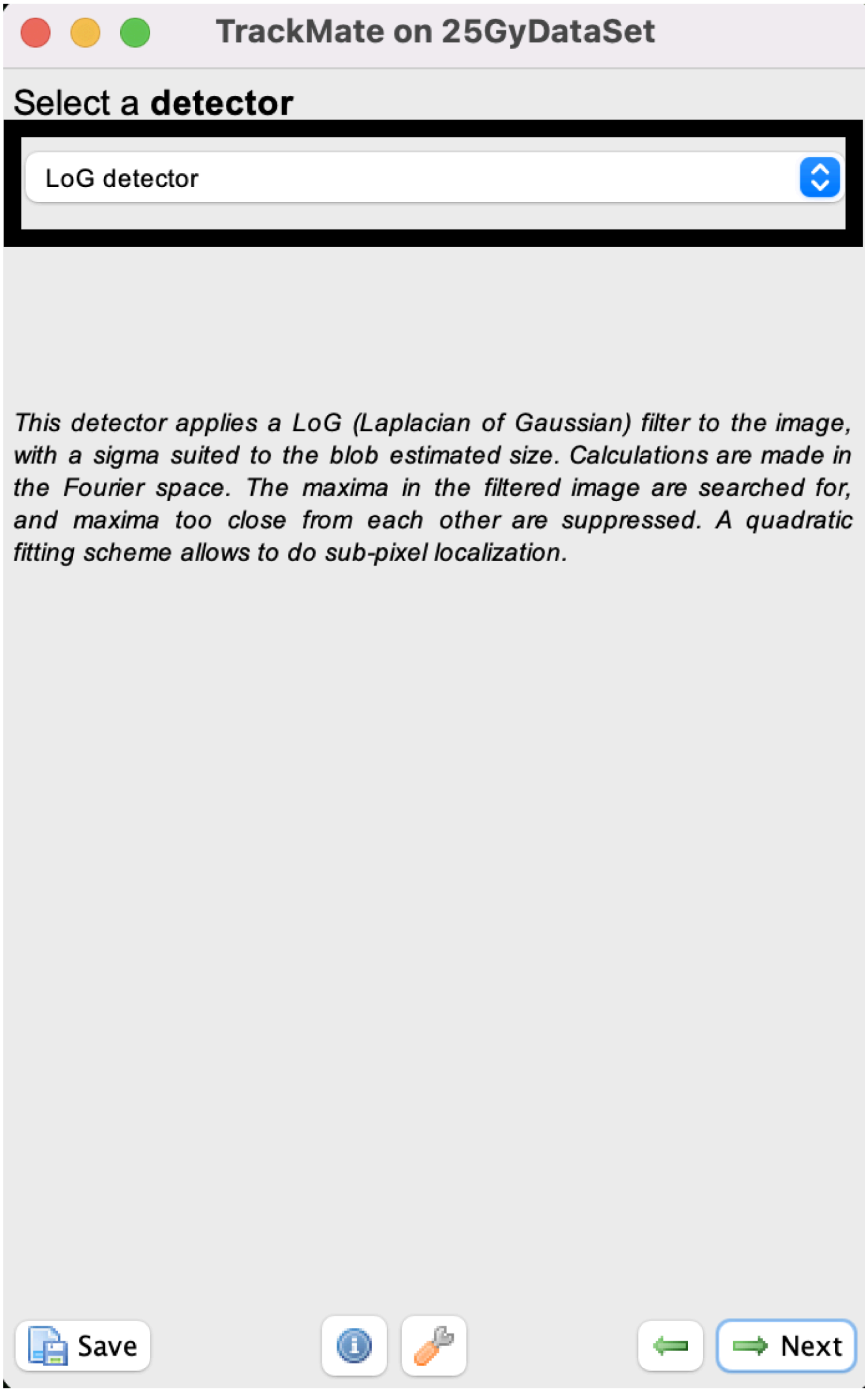


Identify that the LoG detector has been chosen. Click next.

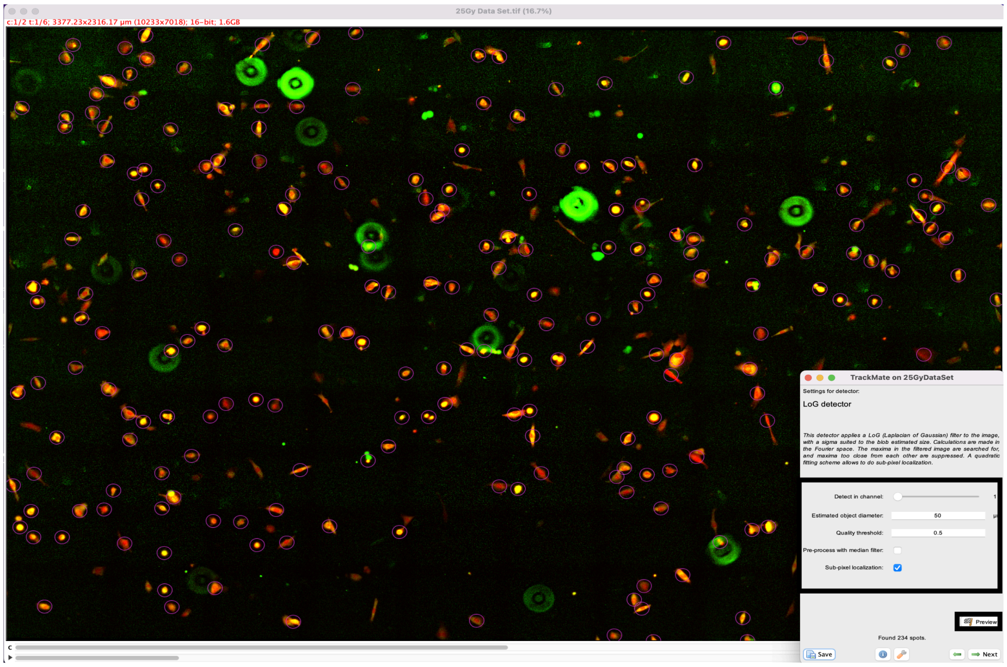


Identify the optimal user-definded tracking settings by previewing the image. Once settings identified click next.

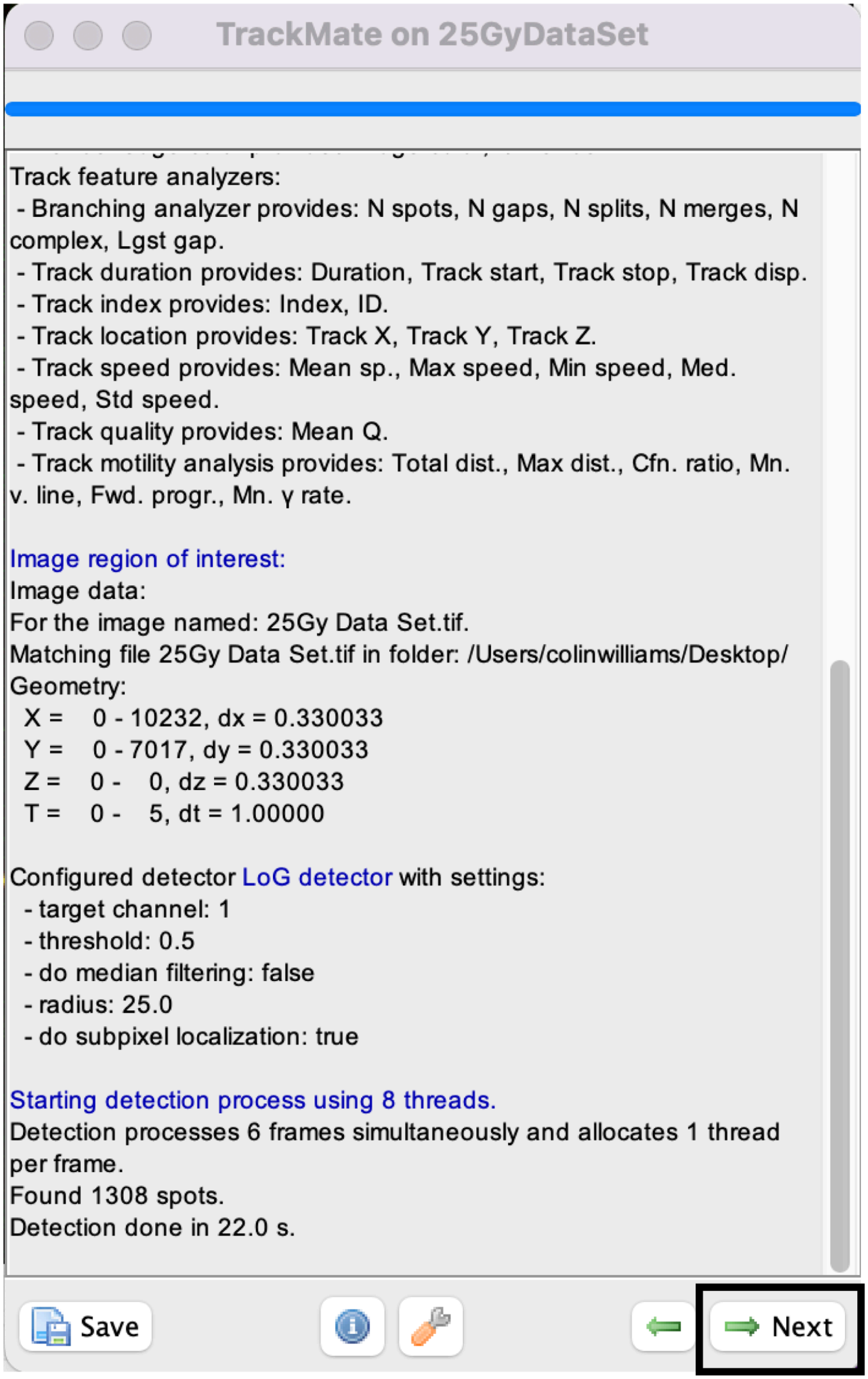


Once detection is complete. Click next.

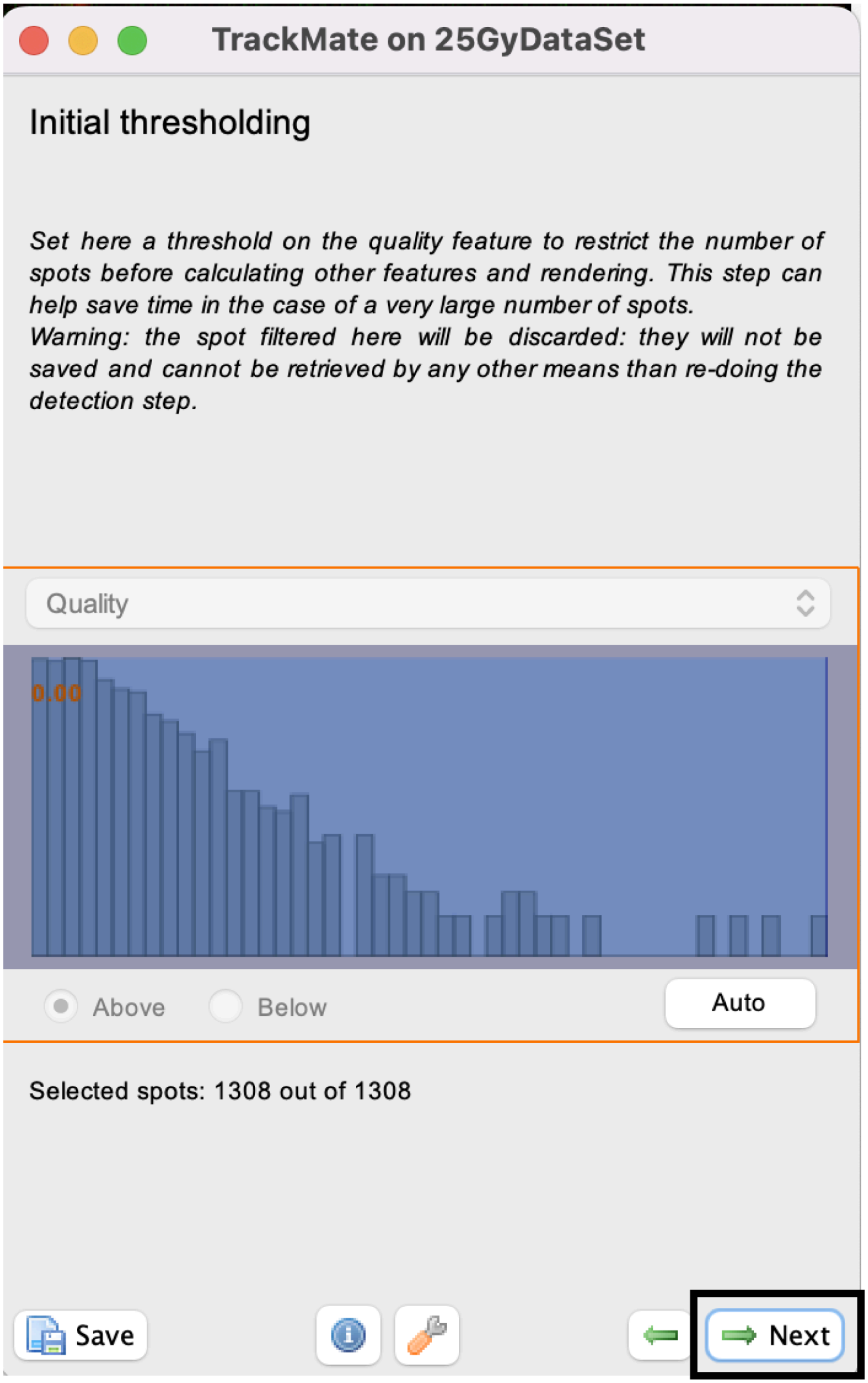


Adjust thresholding of spots if necessary

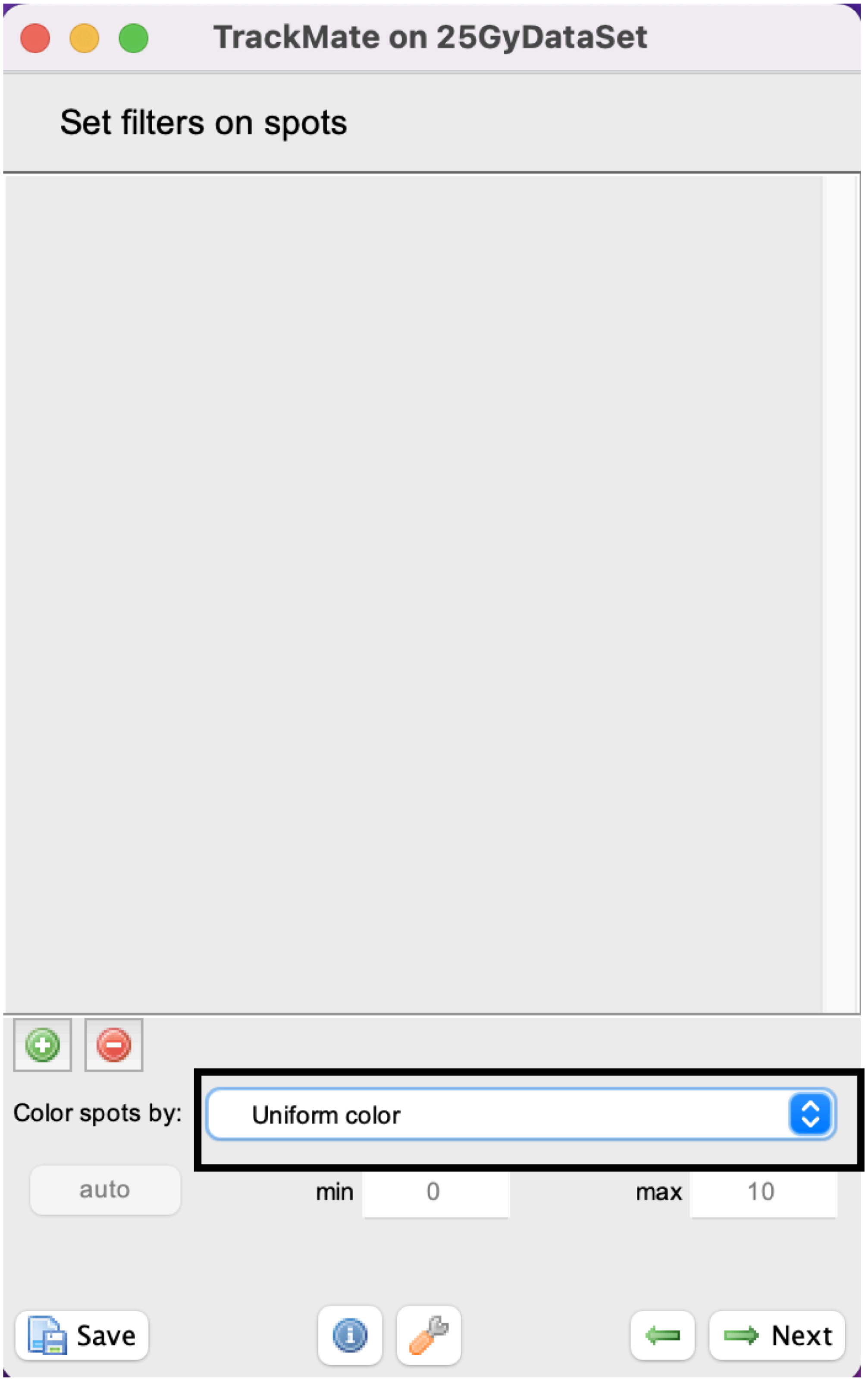


User can set filters to image if there are spots that need to be removed that cannot be removed from adjusting the threshold.

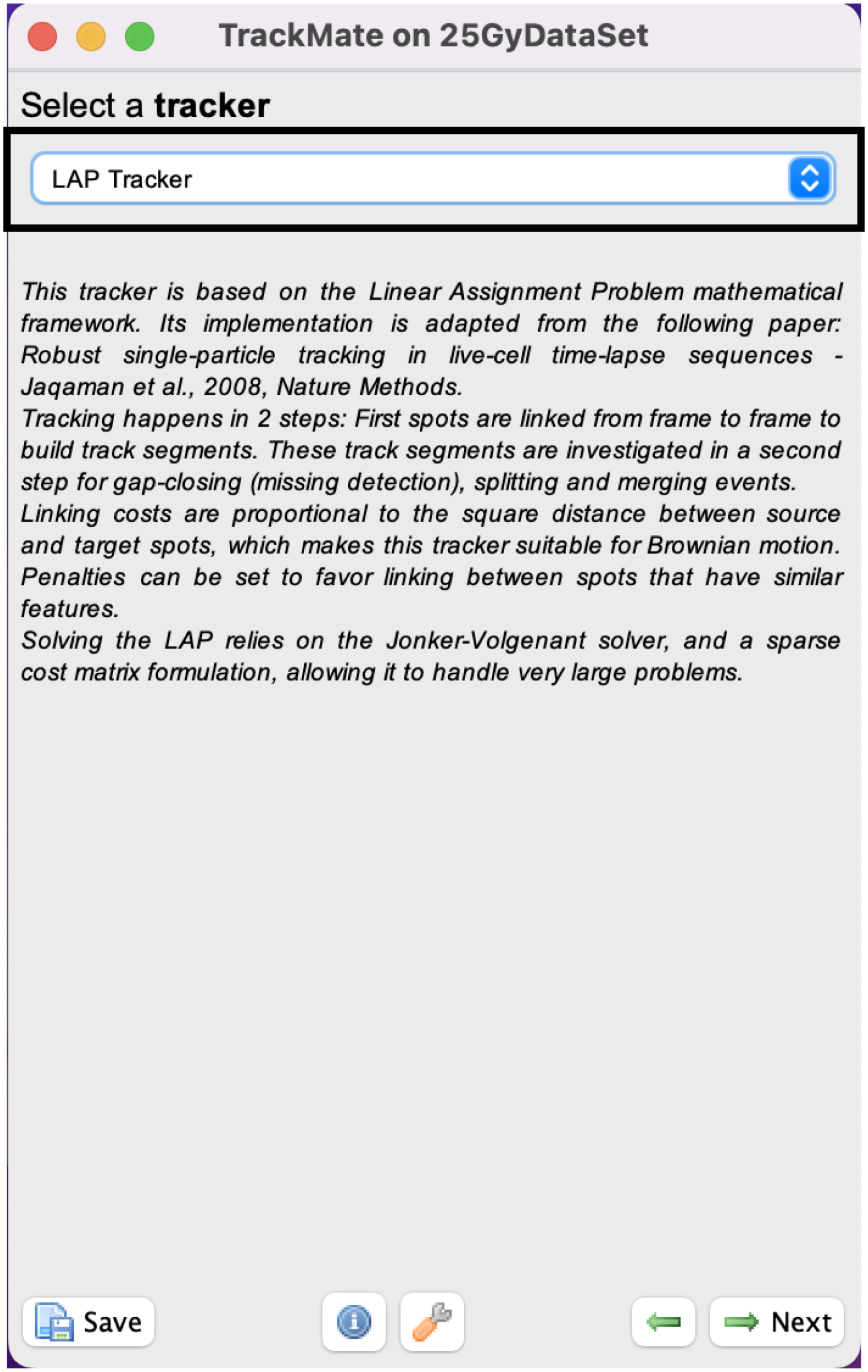


Identify that the LAP tracker has been chosen. Click next.

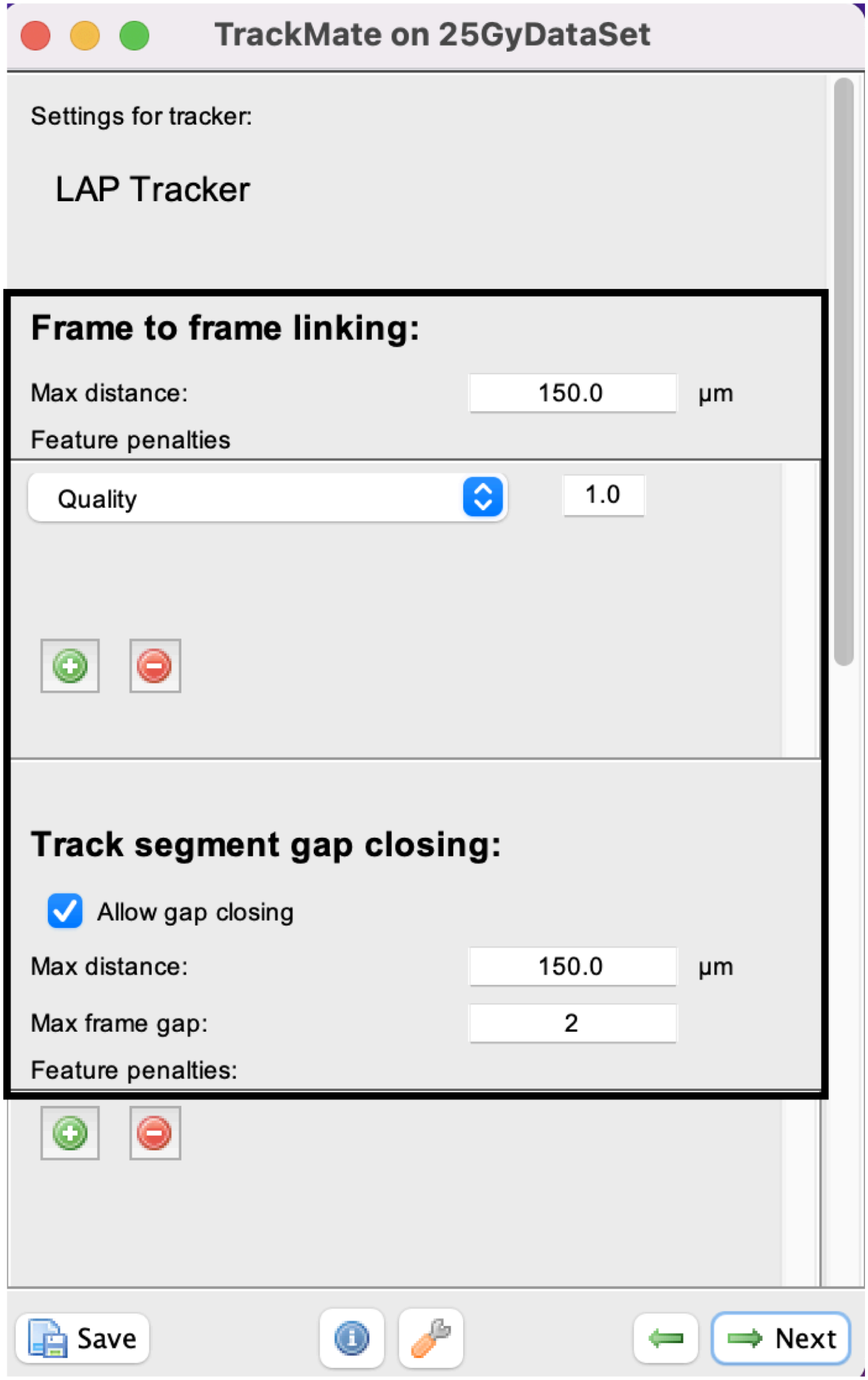


Identify the optimal user-definded tracking settings by previewing the image. The max distance will need iteration to achieve optimal tracking before manual curation.

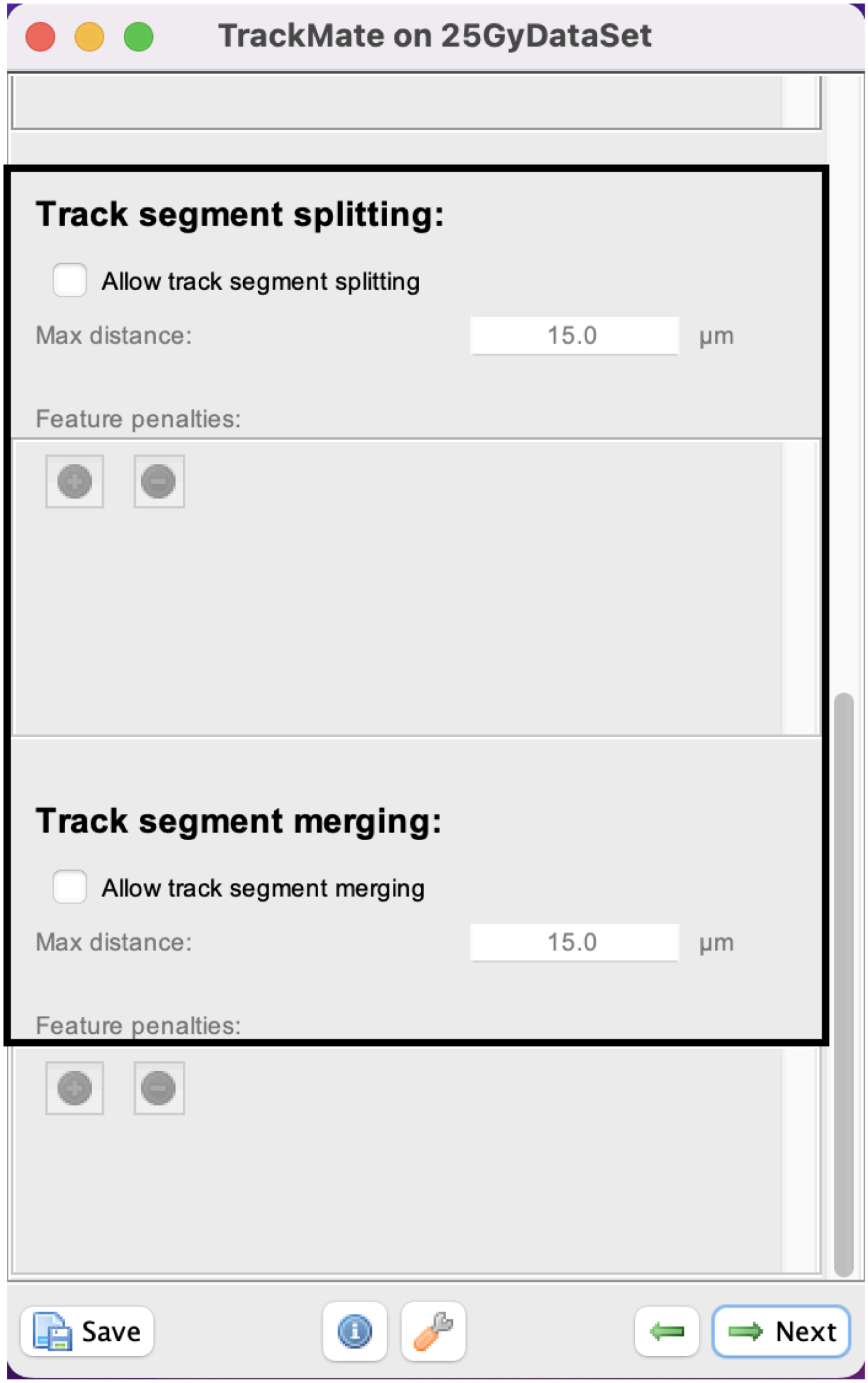


Identify the optimal user-definded tracking settings by previewing the image. Once settings identified click next.

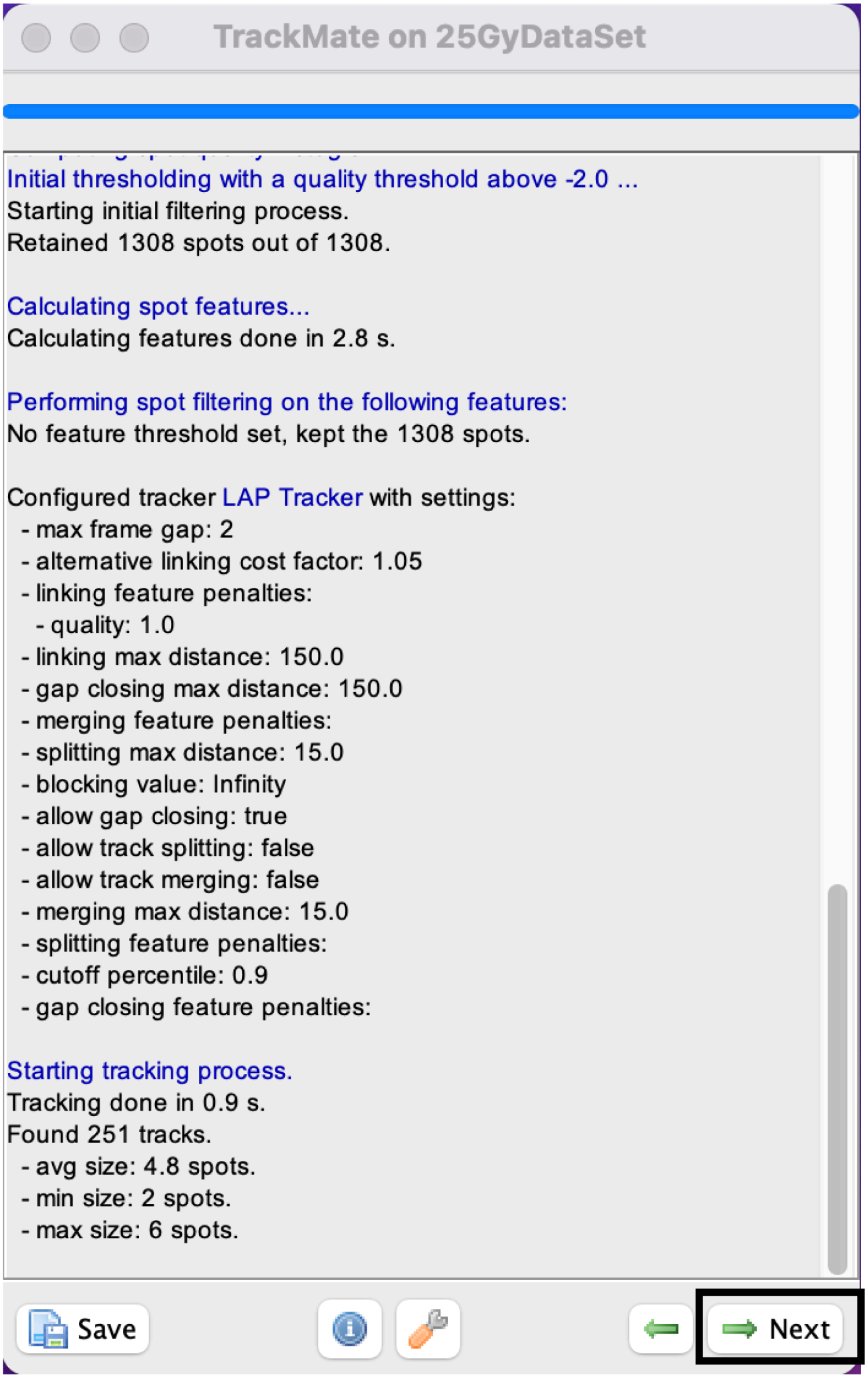


Once detection is complete. Click next.

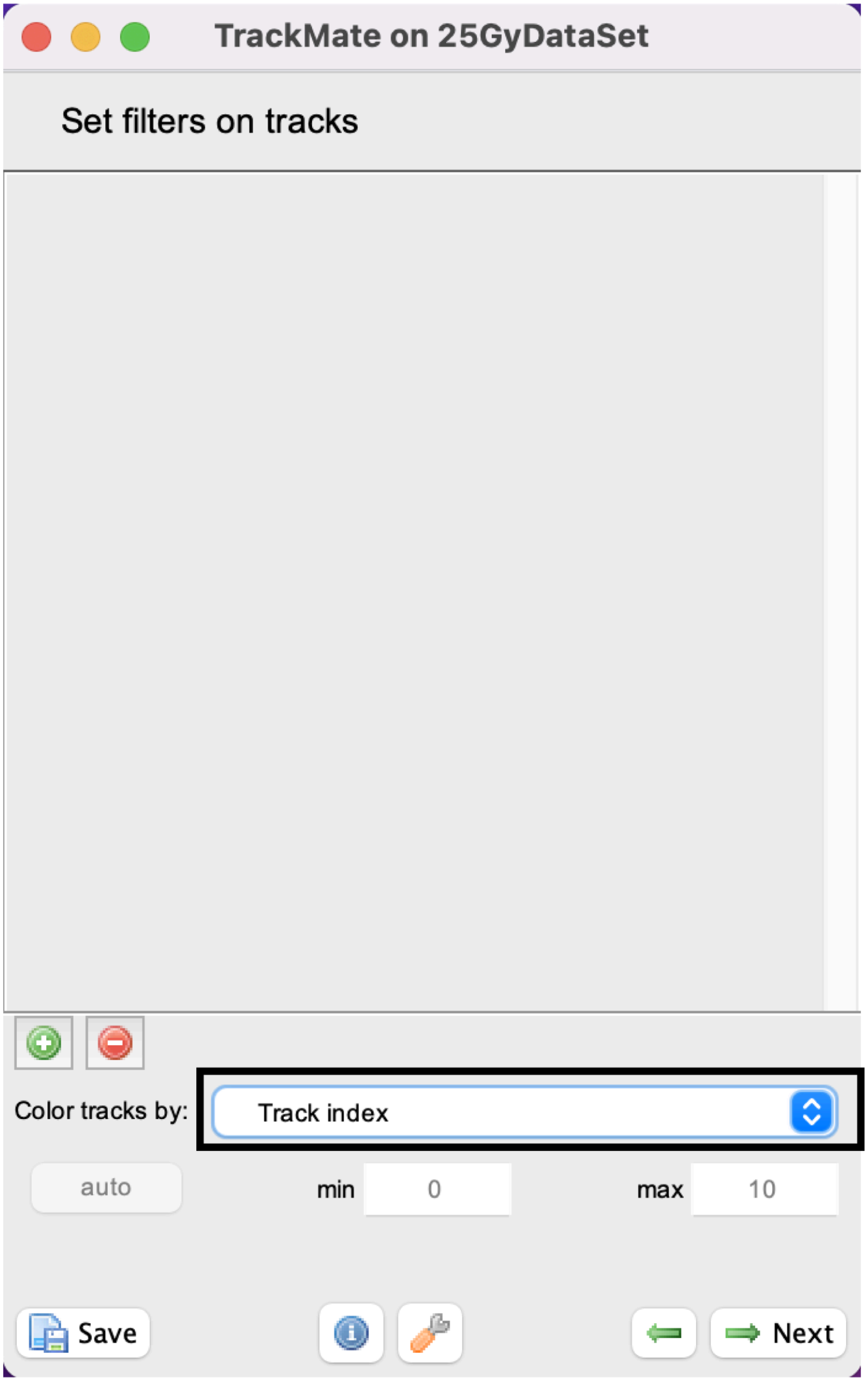


User can set filters to image if tracks need to be removed.

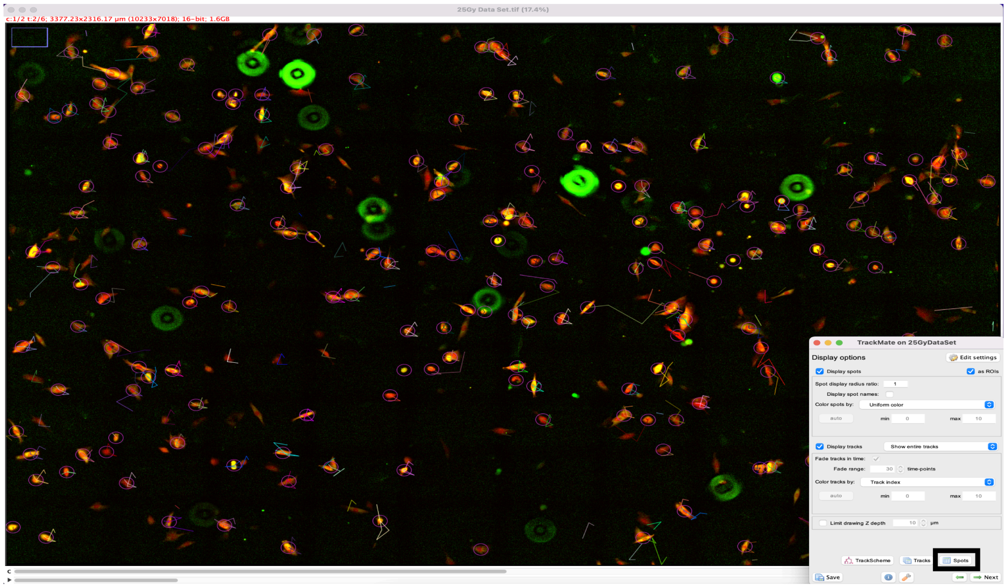


Representative image with spots and tracks.

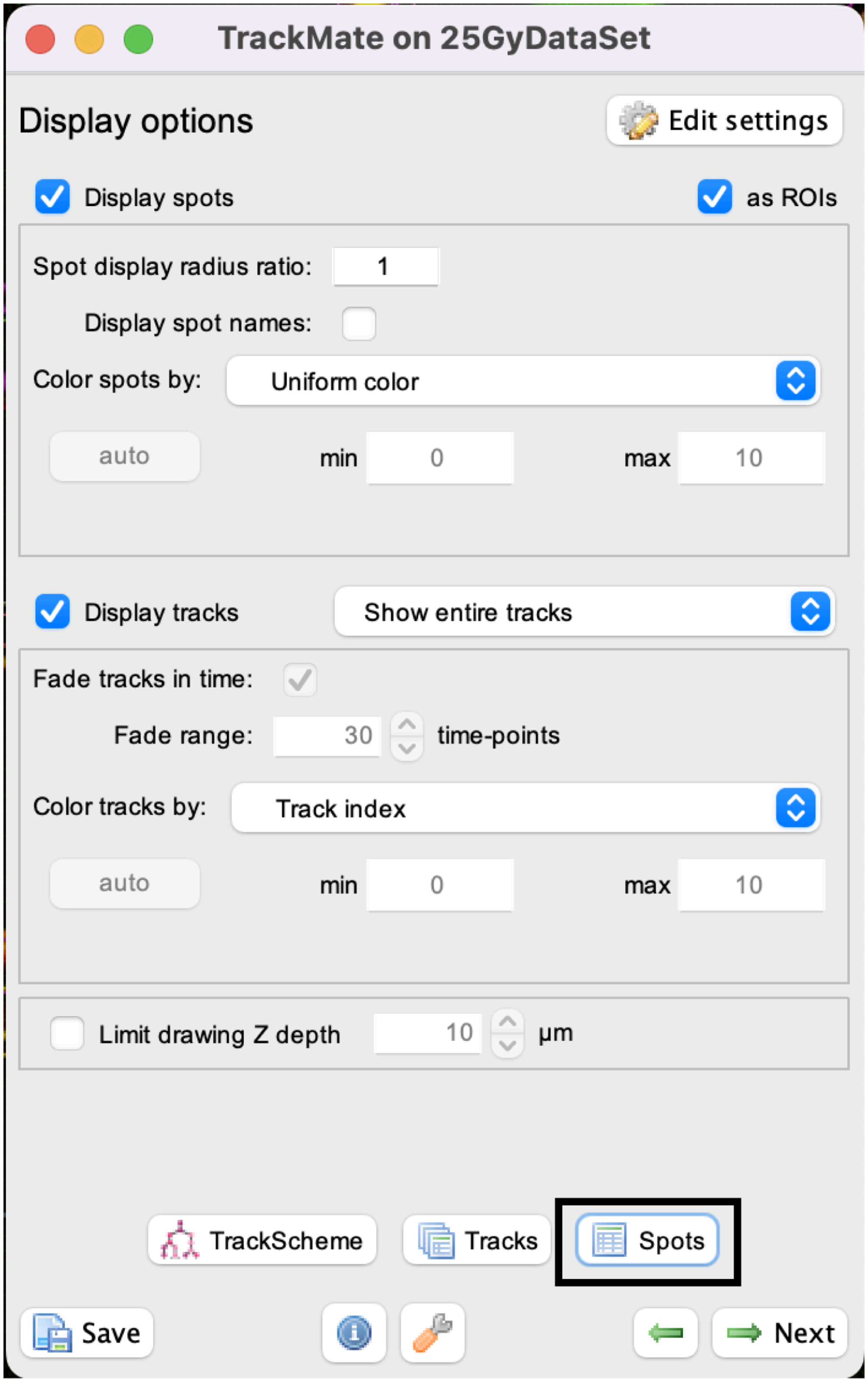


Click the spots button.

All spots table

Export to CSV | Search | coloring

| Label | Spot ID | Track ID | Quality (quality) | X (µm) | Y (µm) | Z (µm) | T (frame) | Frame | R (µm) | Visibility | Spot color | Mean ch1 (counts) |
|---|---|---|---|---|---|---|---|---|---|---|---|---|
| ID4096 | 4096 | 70 | 1.434 | 1,626.259 | 826.999 | 0 | 0 | 0 | 25 | 1 | | 1,484.11 |
| ID4097 | 4097 | | 0.586 | 2,040.932 | 830.532 | 0 | 0 | 0 | 25 | 1 | | 620.743 |
| ID4098 | 4098 | 73 | 1.315 | 2,140.525 | 837.64 | 0 | 0 | 0 | 25 | 1 | | 1,408.39 |
| ID4099 | 4099 | 71 | 0.873 | 1,759.465 | 839.628 | 0 | 0 | 0 | 25 | 1 | | 1,328.615 |
| ID4100 | 4100 | 74 | 1.629 | 913.203 | 847.225 | 0 | 0 | 0 | 25 | 1 | | 1,566.405 |
| ID4101 | 4101 | 75 | 0.788 | 2,711.573 | 847.996 | 0 | 0 | 0 | 25 | 1 | | 1,267.305 |
| ID4102 | 4102 | 72 | 0.819 | 2,320.689 | 851.748 | 0 | 0 | 0 | 25 | 1 | | 1,124.611 |
| ID4103 | 4103 | 76 | 0.965 | 2,871.865 | 852.118 | 0 | 0 | 0 | 25 | 1 | | 1,220.403 |
| ID4104 | 4104 | 206 | 1.222 | 2,595.529 | 874.281 | 0 | 0 | 0 | 25 | 1 | | 1,855.645 |
| ID4105 | 4105 | | 0.603 | 585.076 | 876.348 | 0 | 0 | 0 | 25 | 1 | | 738.163 |
| ID4106 | 4106 | 77 | 0.836 | 2,931.308 | 881.629 | 0 | 0 | 0 | 25 | 1 | | 1,152.501 |
| ID4107 | 4107 | 78 | 1.221 | 973.822 | 888.627 | 0 | 0 | 0 | 25 | 1 | | 2,078.208 |
| ID4108 | 4108 | 211 | 0.867 | 1,667.735 | 908.664 | 0 | 0 | 0 | 25 | 1 | | 1,294.725 |
| ID4109 | 4109 | 79 | 0.659 | 3,002.681 | 920.712 | 0 | 0 | 0 | 25 | 1 | | 871.863 |
| ID4110 | 4110 | 81 | 0.775 | 447.947 | 931.077 | 0 | 0 | 0 | 25 | 1 | | 1,014.075 |
| ID4111 | 4111 | 82 | 1.127 | 3,227.13 | 935.204 | 0 | 0 | 0 | 25 | 1 | | 1,342.415 |
| ID4112 | 4112 | 83 | 1.359 | 2,355.136 | 939.964 | 0 | 0 | 0 | 25 | 1 | | 1,920.985 |
| ID4113 | 4113 | 212 | 1.18 | 2,195.569 | 952.955 | 0 | 0 | 0 | 25 | 1 | | 1,395.821 |
| ID4114 | 4114 | 86 | 1.722 | 1,438.561 | 964.462 | 0 | 0 | 0 | 25 | 1 | | 2,181.914 |
| ID4115 | 4115 | 89 | 1.182 | 2,530.04 | 972.196 | 0 | 0 | 0 | 25 | 1 | | 1,560.434 |
| ID4116 | 4116 | 87 | 1.205 | 1,238.587 | 977.013 | 0 | 0 | 0 | 25 | 1 | | 1,544.263 |
| ID4117 | 4117 | 213 | 0.752 | 1,509.989 | 980.756 | 0 | 0 | 0 | 25 | 1 | | 845.813 |
| ID4118 | 4118 | 84 | 3.851 | 88.563 | 982.149 | 0 | 0 | 0 | 25 | 1 | | 4,178.335 |
| [illegible] | [illegible] | [illegible] | [illegible] | [illegible] | [illegible] | [illegible] | [illegible] | [illegible] | [illegible] | [illegible] | | [illegible] |

Export file as a CSV. The CSV file will be used to identify the GEDI ratio by mean channel 1 divided by mean channel 2. The channel being divided depends on which channel is GC150. For our experiment, GC150 was channel 1.

51 Using annotated tracks and GEDI ratios generate survival curves based off the experimental parameters of interest.

52 Use the intensities quantified by the spot detector to make a new column titled GEDI ratio for the dataset of interest.

53 Ensure each track is validated either manually or using the numerous filters that are available in TrackMate. Type of filters to use will be highly dependent on user generated images.
